# OTULIN protects the liver against cell death, inflammation, fibrosis, and cancer

**DOI:** 10.1101/776021

**Authors:** Rune Busk Damgaard, Helen E. Jolin, Michael E.D. Allison, Susan E. Davies, Andrew N.J. McKenzie, David Komander

**Author notes:** Correspondence (R.B.D); (D.K).

## Abstract

The deubiquitinase OTULIN removes methionine-1 (M1)-linked polyubiquitin chains to regulate TNF-mediated inflammation and cell death, but the physiological role of OTULIN outside the immune system is poorly understood. Here, we identify OTULIN as a liver tumour suppressor in mice. Hepatocyte-specific OTULIN deletion causes spontaneous steatohepatitis, extensive fibrosis, and pre-malignant tumours by eight weeks of age, which progresses to hepatocellular carcinoma by 7-12 months. OTULIN deficiency triggers apoptosis and inflammation in the liver, but surprisingly, steatohepatitis and pre-malignant growth is independent of TNFR1 signalling. Instead, the pathology in OTULIN-deficient livers is associated with increased mTOR activation, and mTOR inhibition with rapamycin reduces fibrosis and pre-malignant growth. This demonstrates that OTULIN is critical for maintaining liver homeostasis and preventing mTOR-driven liver disease.

## Introduction

Liver cancer is the fifth most common type of cancer and the third leading cause of cancer-related deaths worldwide (Llovet et al., 2016). The main risk factors for development of liver cancer are cirrhosis, hepatitis B virus (HBV) or hepatitis C virus (HCV) infection, excessive alcohol consumption, and non-alcoholic steatohepatitis associated with obesity and metabolic syndrome (Llovet et al., 2016). Hepatocellular carcinoma (HCC) is the most common form of liver cancer. Hallmarks of HCC development include chronic inflammation and metabolic alterations, which mechanistically are, at least in part, attributed to hepatocyte cell death and excessive mTOR signalling (Llovet et al., 2016; Luedde and Schwabe, 2011; Mossmann et al., 2018). However, the pathogenesis of HCC remains incompletely understood.

Chronic inflammation caused by persistent damage to the liver parenchyma is a key trigger of HCC development (Luedde et al., 2014). In the context of liver inflammation, sustained compensatory proliferation induced by hepatocyte cell death is pro-tumourigenic and leads to accumulation of mutations and epigenetic changes over time (Liu et al., 2014; Luedde et al., 2014). Pro-inflammatory cytokines and chemokines in the microenvironment support the continuous proliferation and expansion of pre-neoplastic cells, eventually leading to hepatocyte transformation and cancer (Grivennikov et al., 2010). Understanding the cellular processes that contribute to the pathogenesis of chronic liver inflammation resulting in HCC is therefore important to identify new and better therapeutic strategies.

Multiple key regulatory mechanisms in inflammation rely on signalling via non-degradative protein ubiquitination (Hu and Sun, 2016). M1-linked ubiquitin (Ub) chains (hereafter referred to as M1-polyUb) are conjugated by the linear Ub chain assembly complex (LUBAC) and play a central role in regulation of pro-inflammatory nuclear factor *κ*B (NF-*κ*B) signalling, gene activation, and cell death in response to engagement of tumour necrosis factor (TNF) receptor 1 (TNFR1) and a range of other immune receptors (Hrdinka and Gyrd-Hansen, 2017; Shimizu et al., 2015). Upon engagement of TNFR1, LUBAC is recruited to the receptor signalling complex where it conjugates M1-polyUb to facilitate I*κ*B kinase (IKK) and NF-*κ*B activation (Hrdinka and Gyrd-Hansen, 2017; Kupka et al., 2016). However, in the absence of LUBAC and M1-polyUb, TNFR1 signalling is shifted from pro-inflammatory gene activation towards induction of cell death (Brenner et al., 2015; Kupka et al., 2016). Depending on context, this cell death can occur via caspase-dependent apoptosis or caspase-independent necroptosis (Berger et al., 2014; Kumari et al., 2014; Peltzer et al., 2014; 2018; Rickard et al., 2014; Taraborrelli et al., 2018). Dysregulated TNFR1 and NF-*κ*B signalling have been implicated in the pathogenesis of hepatitis and HCC (Luedde and Schwabe, 2011; Sun and Karin, 2008) and several studies have connected regulators of M1-polyUb signalling to the development of liver disease. Both NF-*κ*B essential modulator (NEMO), which is the M1-polyUb-selective receptor required for IKK activation (Rahighi et al., 2009), and HOIL-1-interacting protein (HOIP), the catalytic subunit of LUBAC, protect against liver damage and carcinogenesis by preventing hepatocyte apoptosis and inflammation (Kondylis et al., 2015; Luedde et al., 2007; Shimizu et al., 2017). This indicates that proper regulation of M1-polyUb signalling is a checkpoint in liver homeostasis.

Ubiquitin signalling is antagonised by deubiquitinases (DUBs), which cleave the polyUb signal from substrates to terminate signalling (Clague et al., 2019). Multiple DUBs are implicated in cancer development. They can act as both oncogenes and tumour suppressors (Fraile et al., 2011), and DUBs have become attractive drug targets for development of new anti-cancer therapeutics (Harrigan et al., 2018). The DUBs CYLD and A20, which regulate NF-*κ*B activation and cell death downstream of TNFR1 and other immune receptors (Lork et al., 2017), act as tumour suppressors and are mutated in a range of cancers (Sun, 2010; Zhang et al., 2012). In mice, both CYLD and A20 protect against hepatitis and HCC development by preventing apoptosis of hepatocytes (Catrysse et al., 2016; Luedde et al., 2007).

OTU family deubiquitinase with linear Ub specificity (OTULIN) and CYLD are the two main DUBs that regulate M1-polyUb signalling (Elliott, 2016; Lork et al., 2017). OTULIN exclusively cleaves M1 linkages (Keusekotten et al., 2013; Rivkin et al., 2013), whereas CYLD cleaves both K63 and M1 linkages (Komander et al., 2008). OTULIN binds directly to the LUBAC subunit HOIP (Elliott et al., 2014; Schaeffer et al., 2014; Takiuchi et al., 2014) and regulates LUBAC signalling, autoubiquitination, and stability (Damgaard et al., 2016; 2019; Fiil et al., 2013; Heger et al., 2018; Hrdinka et al., 2016; Keusekotten et al., 2013). In humans, homozygous hypomorphic mutations in *OTULIN* cause OTULIN-related autoinflammatory syndrome (ORAS) (also known as otulipenia, OMIM #617099), a life-threatening autoinflammatory disease characterised by sterile systemic inflammation, recurrent high fevers, panniculitis, diarrhoea, arthritis, and general failure to thrive (Damgaard et al., 2016; 2019; Nabavi et al., 2019; Zhou et al., 2016). The primary driver of inflammation in OTULIN-deficient humans and mice is TNF signalling (Damgaard et al., 2016; Zhou et al., 2016), which in myeloid cells leads to LUBAC hyper-signalling and NF-*κ*B activation (Damgaard et al., 2016; 2019). In other cell types, e.g. fibroblasts, OTULIN loss leads to LUBAC degradation and TNF-induced cell death (Damgaard et al., 2019; Heger et al., 2018). However, it remains unclear if OTULIN deficiency may cause or predispose to development of other pathologies, for example cancer.

Here we identify OTULIN as a critical factor for maintaining liver homeostasis. Hepatocyte-specific genetic ablation of OTULIN in mice led to spontaneous steatosis, hepatitis, and fibrosis. This triggered tumour formation by the age of eight weeks, resulting in development of HCC by 7-12 months. Surprisingly, the pathology in OTULIN-deficient livers was independent of TNFR1-mediated signalling. Instead, early tumourigenesis in these mice was partially dependent on mTOR signalling, and treatment with the mTOR inhibitor rapamycin reduced liver pathology. Our results identify OTULIN as a tumour suppressor in the liver and show that OTULIN and correct regulation of M1-polyUb signalling are crucial for preventing spontaneous and severe liver disease.

## Results

### *Otulin* deletion in non-haematopoietic cells causes acute hepatitis and liver failure

Mice with germline mutation or inactivation of OTULIN die *in utero*. Embryos carrying two different hypomorphic OTULIN mutations (W96R or D336E) die between embryonic day (E)12.5-E14, attributed to impaired Wnt signalling (Rivkin et al., 2013). Expression of an active site cysteine mutant (C129A), which renders OTULIN catalytically inactive but converts it to a high-affinity M1-polyUb-binding protein (Fiil et al., 2013; Keusekotten et al., 2013; Morrow et al., 2018), leads to embryonic lethality at E10.5 due to excessive caspase-8- and RIPK3-dependent cell death (Heger et al., 2018). Generation of bone marrow chimeras and conditional *Otulin* knockout (KO) mice have identified cell type-specific phenotypes of OTULIN-deficiency in immune cells in adult mice (Damgaard et al., 2016), but the role of OTULIN in most non-haematopoietic cell types in adult mice is unknown. To investigate the physiological role of OTULIN specifically in non-haematopoietic cells, we replaced the bone marrow of *Rosa26*-Cre-ERT2-*Otulin*^flox^ mice (Damgaard et al., 2016) with wild type bone marrow to generate chimeric mice that become OTULIN-deficient in all non-haematopoietic cells after tamoxifen administration (*Otulin*-KO^Chim^ mice) (**Figure 1A**).

**Figure 1.**
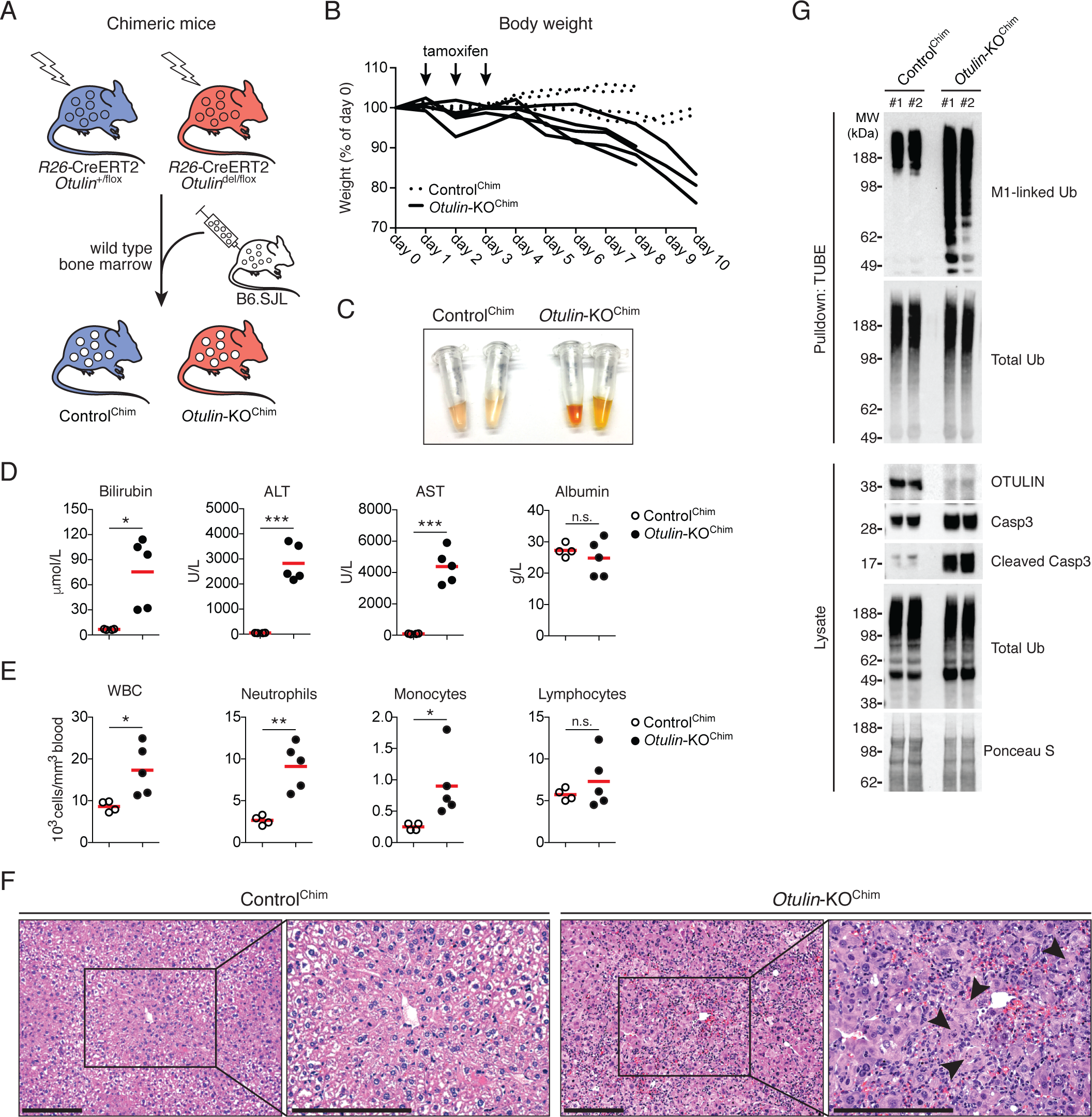
Deletion of OTULIN in non-haematopoietic cells causes rapid liver failure. **(A)** Schematic representation of wild type B6.SJL bone marrow transplantation into *γ*-irradiated *Rosa26*-Cre-ERT2-*Otulin* (*R26*-CreERT2-*Otulin*) mice. **(B)** Relative body weight following i.p. administration of tamoxifen (arrows) to Control^Chim^ (n=4) and *Otulin*-KO^Chim^ (n=5) mice. Each line represents one mouse. Data were pooled from two independent experiments. **(C)** Serum from terminal bleeds of Control^Chim^ and *Otulin*-KO^Chim^ mice at the end of the experiment shown in (B). **(D-E)** Analysis of bilirubin, ALT, AST, and albumin levels in serum (D) and blood cell counts (E) from terminal bleeds of Control^Chim^ (n=4) and *Otulin*-KO^Chim^ (n=5) mice at the end of the experiment shown in (B). Data were pooled from two independent experiments. Data are presented as individual data points, each representing one mouse. Red bars indicate means. Data were analysed using an unpaired, two-sided Student’s *t* test. n.s., non-significant. **(F)** Micrographs of H&E stained liver sections from Control^Chim^ and *Otulin*-KO^Chim^ mice at the end of the experiments shown in (B). Arrowheads indicate cells with nuclear condensation and fragmentation. Micrographs are representative of two mice in each group. **(G)** Immunoblot analysis of whole-liver lysates and endogenous Ub conjugates purified by TUBE pulldown from livers of two Control^Chim^ and two *Otulin*-KO^Chim^ mice at the end of the experiment shown in (B). See also Figure S1.

Six weeks after transplantation, the chimeric mice were treated with tamoxifen to induce *Otulin* deletion. This resulted in 10-20% weight loss over 8-10 days in the *Otulin*-KO^Chim^ mice (**Figure 1B)**. Weight loss was accompanied by highly icteric serum (**Figure 1C**) with a ∼12-fold increase in the serum level of the haem metabolite bilirubin (**Figure 1D**), indicating potential liver failure in *Otulin*-KO^Chim^ mice. The liver enzymes alanine aminotransferase (ALT) and aspartate aminotransferase (AST) were also strongly increased in the *Otulin*-KO^Chim^ serum (**Figure 1D**), indicating damage to the liver parenchyma, and the number of circulating white blood cells, particularly neutrophils, were elevated in the blood (**Figure 1E**). Histological analysis confirmed severe acute hepatitis in the *Otulin*-KO^Chim^ mice with immune cell infiltration and multiple dead or dying hepatocytes with nuclear condensation and fragmentation in the liver (**Figure 1F**). In contrast, we observed no obvious pathology in the small intestine, colon, kidney, skin, lung, heart, skeletal muscle, or adipose tissue when compared with Control^Chim^ mice (**Figure S1A**). Immunoblot analysis confirmed efficient deletion of OTULIN in the *Otulin*-KO^Chim^ livers (**Figures 1G and S1B**), and tandem Ub-binding entity (TUBE)-mediated enrichment of Ub conjugates showed an increase of M1-polyUb in these livers compared with controls, confirming deregulated M1-polyUb conjugation in the *Otulin*-KO^Chim^ mice. Strikingly, OTULIN deficiency led to marked cleavage and activation of caspase-3 (**Figure 1G**), suggesting that the liver pathology in *Otulin*-KO^Chim^ mice could be driven by apoptosis. Only ∼2% of CD45^+^ immune cells present in peripheral tissues in the chimeric mice were of parental origin (**Figures S1C-D**), indicating that the observed phenotype is caused by intrinsic defects in the hepatocytes with minimal contribution from OTULIN-deficient immune cells. Collectively, these results show that OTULIN has important physiological functions in non-haematopoietic cells and is critical for preventing hepatocyte damage and hepatitis.

### Hepatocyte-specific loss of OTULIN causes spontaneous steatohepatitis, fibrosis, and tumourigenesis

To study the role of OTULIN and M1-polyUb signalling in the liver specifically, we generated mice with hepatocyte-specific deletion of OTULIN by breeding mice carrying the *Otulin*^flox^ allele (Damgaard et al., 2016) with transgenic mice expressing the Cre recombinase under the control of a serum albumin promoter (*Alb*-Cre) (Postic et al., 1999) (**Figure S2A**). Mice with *Otulin* deletion in the liver (*Otulin*^Δhep^ mice) were born and weaned at the expected Mendelian frequency but developed obvious liver pathology (**Figure 2A**). OTULIN protein levels were efficiently reduced in whole-liver lysates from these livers (**Figures 2B and S2B**). Similar to the *Otulin*-KO^Chim^ mice, OTULIN loss caused a concomitant increase in the level of M1-polyUb in the *Otulin*^Δhep^ livers compared with controls (**Figures 2C and S2C**), confirming deregulated M1-polyUb signalling. Residual expression of OTULIN in *Otulin*^Δhep^ livers can be attributed to incomplete penetrance of *Alb*-Cre-mediated gene deletion in hepatocytes (**Figures 2B and S2D**) as well as to non-parenchymal liver cells that are not targeted by *Alb*-Cre. Expression of the LUBAC components HOIP, HOIL-1, and SHARPIN was reduced, similar to the effect of OTULIN deficiency observed in lymphocytes or fibroblasts (Damgaard et al., 2016; 2019), while the level of CYLD remained unchanged (**Figure 2B**).

**Figure 2.**
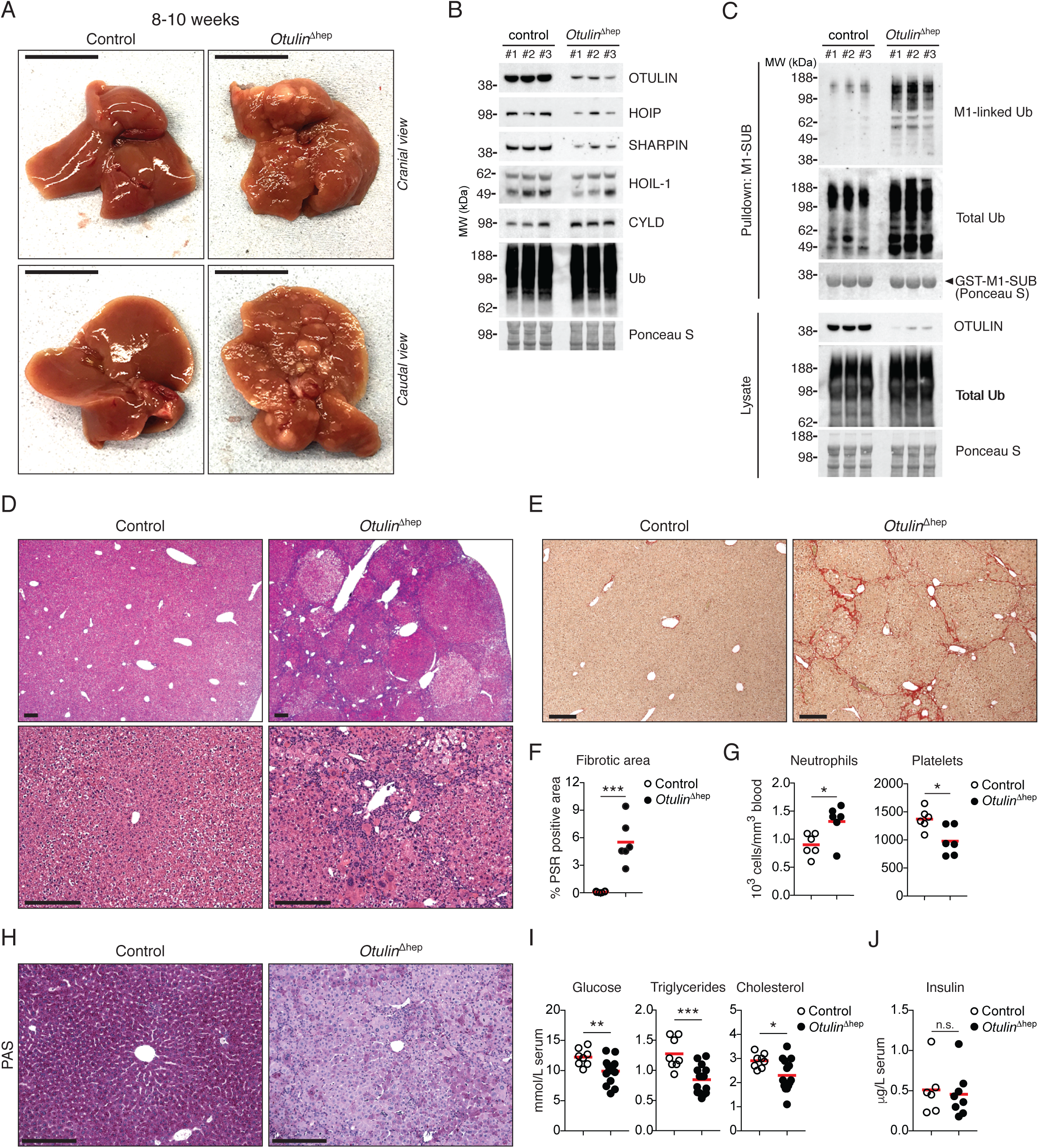
Steatohepatitis, fibrosis, and spontaneous tumour formation in *Otulin*^Δhep^ mice. **(A)** Representative macroscopic appearance of *Otulin*^Δhep^ and control livers at the age of 8-10 weeks. Scale bars indicate 1 cm. **(B)** Immunoblot analysis of OTULIN, LUBAC, and CYLD expression in whole-liver lysates from three *Otulin*^Δhep^ and three control mice aged 8-10 weeks. **(C)** Immunoblot analysis of whole-liver lysates and endogenous Ub conjugates purified by M1-SUB pulldown from livers of three control and three *Otulin*^Δhep^ mice. **(D)** Micrographs of H&E stained liver sections from *Otulin*^Δhep^ and control mice aged 8-10 weeks. Top panels show pale-staining hepatocyte clones with fat accumulation in *Otulin*^Δhep^ mice. Bottom panels show inflammation, fat accumulation, and variations in nuclear size in *Otulin*^Δhep^ livers. Micrographs are representative of eight mice of each genotype. **(E)** Micrographs of PSR stained liver sections from *Otulin*^Δhep^ and control mice aged 8-10 weeks show fine bridging porto-portal and porto-central fibrous septa with areas of pericelluar fibrosis in *Otulin*^Δhep^ mice. Micrographs are representative of six mice of each genotype. **(F)** Quantification of PSR-positive (fibrotic) area in liver sections from *Otulin*^Δhep^ (n=6) and control (n=6) mice aged 8-10 weeks. **(G)** Neutrophil and platelet counts from terminal bleeds of *Otulin*^Δhep^ (n=6) and control (n=6) mice aged 8-10 weeks. **(H)** Micrographs of PAS stained liver sections from *Otulin*^Δhep^ and control mice aged 8-10 weeks show pale-staining hepatocytes in *Otulin*^Δhep^ mice due to loss of glycogen. Micrographs are representative of five controls and six *Otulin*^Δhep^ mice of each genotype. **(I-J)** Analysis of glucose, triglyceride, and cholesterol levels in serum from terminal bleeds of *Otulin*^Δhep^ (n=15) and control (n=8) mice aged 8-10 weeks. **(J)** Analysis of insulin levels in serum from terminal bleeds of *Otulin*^Δhep^ (n=8) and control (n=6) mice aged 8-10 weeks. **(F-G, I-J)** Data are presented as individual data points, each representing one mouse. Red bars indicate means. Data were analysed using an unpaired, two-sided Student’s *t* test. n.s., non-significant. See also Figure S2.

Dissection of livers from *Otulin*^Δhep^ mice at the age of 8-10 weeks revealed severe liver disease with multiple macroscopic lesions and nodules present in young adult mice (**Figure 2A**). Microscopic examination showed markedly abnormal liver histology in the *Otulin*^Δhep^ liver parenchyma (**Figure 2D**). Histopathologically, we observed focal steatosis, Mallory-Denk bodies, Kupffer cell hyperplasia, and inflammatory foci in the OTULIN-deficient livers (**Figures 2D and S2E**). These alterations are hallmarks of chronic liver disease with non-alcoholic steatohepatitis (NASH) (Farrell and Larter, 2006). Consistent with NASH-like disease, Picro Sirius Red (PSR) staining showed extensive collagen deposition in the *Otulin*^Δhep^ livers (**Figures 2E-F**), with bridging septa and pericellular fibrosis (**Figure S2G**), resembling the fibrotic lesions in NASH and cirrhosis in humans (Farrell and Larter, 2006). NASH is a risk factor for HCC development (Llovet et al., 2016). Further examination of the *Otulin*^Δhep^ livers confirmed that many of the tumours and nodules observed macroscopically (**Figure 2A**) were in fact dysplastic nodules (**Figures 2D and S2F**). Across the parenchyma, we observed prominent variation in size of nuclei (anisokaryosis), large cell change (LCC), and clone-like growth (**Figures 2D and S2E-F**), which are well-known pre-malignant changes (Libbrecht et al., 2005). This liver pathology was fully penetrant in all *Otulin*^Δhep^ mice analysed at this age, while none of the littermate controls showed any signs of pathology. We therefore conclude that OTULIN is intrinsically important in hepatocytes for preventing severe liver disease.

Despite the absence of hepatomegaly (**Figure S2H**), *Otulin*^Δhep^ mice exhibited multiple additional indications of liver disease. *Otulin*^Δhep^ mice had increased neutrophil counts and decreased platelet counts (**Figure 2G**) in addition to a significant proportion of hepatocytes with polyploid nuclei with a DNA content of 8n or above (**Figures S2I-L**), consistent with findings in cirrhotic livers and NASH (Farrell and Larter, 2006; Gentric et al., 2015; Xu et al., 2014). Intriguingly, *Otulin*^Δhep^ livers also had severely reduced glycogen content. PAS staining, which labels polysaccharides, was homogenous and strong across the tissue in control livers, whereas OTULIN-deficient livers showed weak and heterogenous staining with only a minority of hepatocytes containing diffuse PAS-positive inclusions (**Figure 2H**). The reduced glycogen content was associated with decreased serum levels of glucose, triglycerides, and cholesterol (**Figure 2I**), despite normal insulin levels (**Figure 2J**). These results indicate a disturbance in metabolic function, which could contribute to development or progression of liver disease in *Otulin*^Δhep^ mice.

### OTULIN deficiency in the liver leads to cell death and inflammation

Hepatocyte damage and cell death promotes inflammation and development of NASH (Luedde et al., 2014). We therefore investigated if the pathology observed in *Otulin*^Δhep^ mice was associated with cell death and inflammation. Compared with controls, we observed more TUNEL-positive dead cells in the OTULIN-deficient livers as well as highly increased numbers of proliferating cells with Ki67-positive nuclei (**Figures 3A-C**). Serum from *Otulin*^Δhep^ mice also contained higher levels of ALT, AST, and bilirubin (**Figures 3D-C**), consistent with hepatocyte cell death and a moderate reduction in liver function, while albumin levels remained normal (**Figure 3E**). Similar to the chimeric mice, immunoblot analysis of liver lysates showed increased caspase-3 activation in *Otulin*^Δhep^ mice compared with controls (**Figures 3F and S3A**).

**Figure 3.**
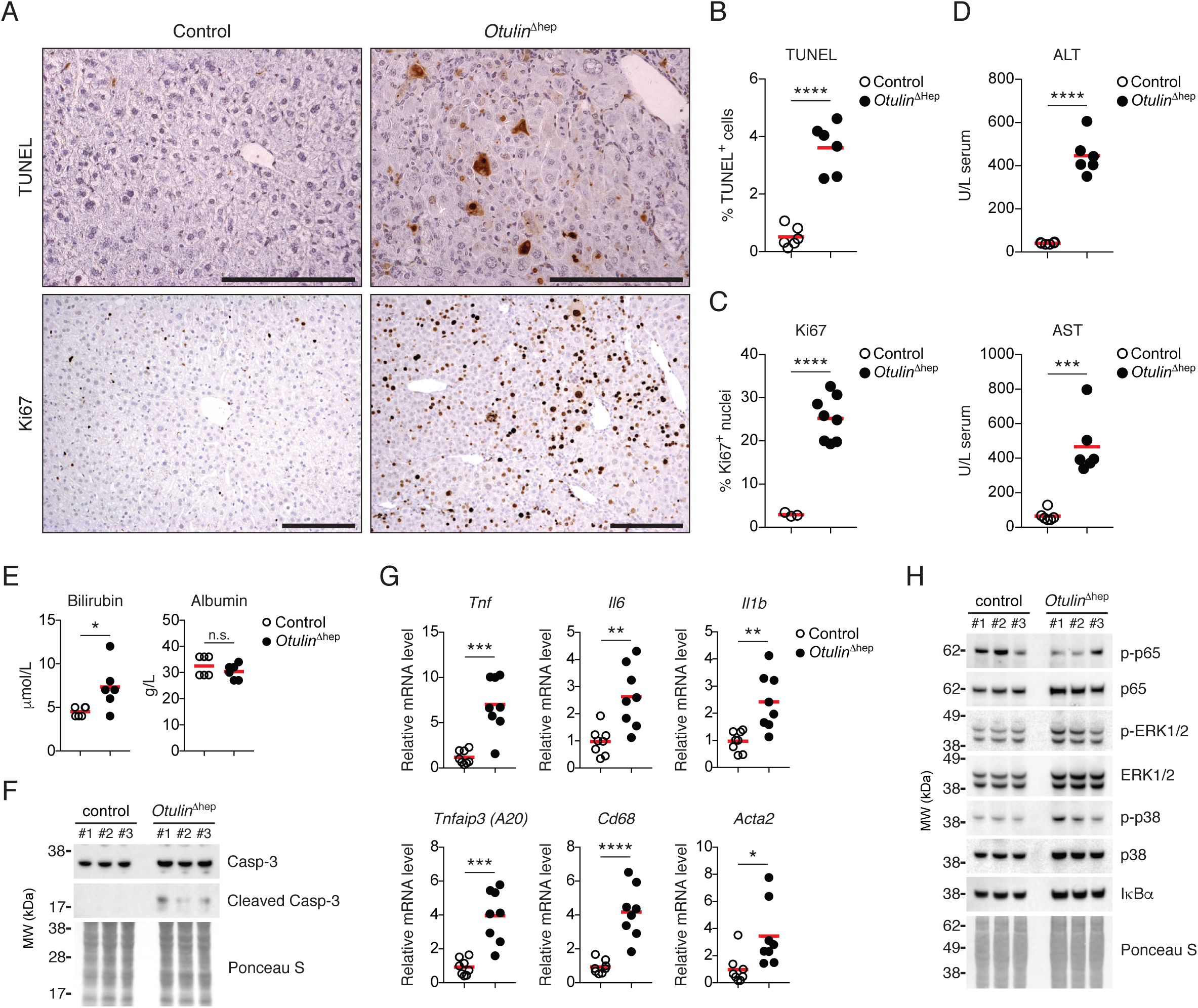
Liver disease in *Otulin*^Δhep^ mice is associated with hepatocyte cell death, proliferation, and inflammation. **(A)** TUNEL (top panels) and anti-Ki67 (bottom panels) stainings of liver sections from *Otulin*^Δhep^ and control mice aged 8-10 weeks. Data are representative of six mice of each genotype for TUNEL staining and three controls and eight *Otulin*^Δhep^ mice for Ki67. **(B-C)** Quantification of TUNEL-(B) and Ki67-positive (C) cells in liver from *Otulin*^Δhep^ and control at the age of 8-10 weeks as shown in (A). TUNEL (B), *Otulin*^Δhep^ (n=6) and control (n=6), and anti-Ki67 (C), *Otulin*^Δhep^ (n=8) and control (n=3). **(D-E)** Analysis of ALT and AST (D) or bilirubin and albumin (E) levels in serum from terminal bleeds of *Otulin*^Δhep^ (n=6) and control (n=6) mice aged 8-10 weeks. **(F)** Immunoblot analysis of caspase-3 cleavage in whole-liver lysate from livers of three control and three *Otulin*^Δhep^ mice aged 8-10 weeks. **(G)** Relative mRNA expression of *Tnf*, *Il6*, *Il1b*, *Tnfaip3*, *Cd68*, and *Acta2* in livers from *Otulin*^Δhep^ (n=8) and control (n=8) aged 8-10 weeks measured by quantitative RT-PCR. **(H)** Immunoblot analysis of NF-*κ*B p65/RelA and MAP kinase activation in whole-liver lysate from livers of three control and three *Otulin*^Δhep^ mice aged 8-10 weeks. **(B-D, E, G)** Data are presented as individual data points, each representing one mouse. Red bars indicate means. Data were analysed using an unpaired, two-sided Student’s *t* test. n.s., non-significant. See also Figure S3.

Cell death and proliferation in the *Otulin*^Δhep^ livers was associated with elevated transcription of inflammatory markers. The mRNA levels of the pro-inflammatory cytokines TNF, IL-6, and IL-1*β* were significantly increased, as were the levels for the NF-*κ*B and apoptosis regulator A20 (*Tnfaip3*) and the Kupffer cell marker CD68 (**Figure 3G**). Cell death and inflammation are key activators of hepatic stellate cell differentiation into collagen-producing myofibroblasts (Luedde and Schwabe, 2011). Consistent with the collagen deposition in OTULIN-deficient livers (**Figure 2D**), the transcript level of smooth muscle actin (*Acta2*), a myofibroblast marker, was also significantly increased (**Figure 3G**), implying expansion of myofibroblasts in the *Otulin*^Δhep^ livers. Interestingly, the inflammation in the OTULIN-deficient livers was not associated with any appreciable increase in NF-*κ*B or MAP kinase activation, at least not at the level of a whole-liver lysate. Immunoblot analysis showed that phosphorylation and activation of the NF-*κ*B family member p65/RelA and the MAP kinases ERK1/2 and p38 was similar in *Otulin*^Δhep^ and control livers, as were the levels of the NF-*κ*B inhibitor Inhibitor-of-*κ*B*α* (I*κ*B*α*) (**Figures 3H and S3B**). The absence of increased inflammatory signalling is similar to previous reports from OTULIN-deficient fibroblasts, which are sensitised to induction of apoptosis rather than NF-*κ*B hyper-signalling (Damgaard et al., 2019; Heger et al., 2018), suggesting that apoptotic cell death of hepatocytes could drive the progressively expanding fibrotic lesions and NASH-like liver pathology observed in *Otulin*^Δhep^ mice.

### Development of hepatocellular carcinoma in OTULIN-deficient livers

Chronic inflammation, NASH, and cirrhosis predispose to development of HCC (Llovet et al., 2016; Luedde and Schwabe, 2011). To examine if the NASH-like pathology in the young *Otulin*^Δhep^ mice might lead to cancer, we analysed the OTULIN-deficient livers for signs of neoplasia and HCC. *Otulin*^Δhep^ mice aged 8-10 weeks had multiple pre-malignant tumours (**Figures 4A and 2D**) and ∼60 macroscopic lesions per liver (**Figure 4B**). The presence of pre-malignant lesions was accompanied by a dramatic increase in serum levels of the liver cancer marker Alpha-Fetoprotein (AFP) (**Figure 4C**). In addition, many genes associated with HCC development were significantly increased in expression in *Otulin*^Δhep^ livers already at 8-10 weeks, including the HCC markers *Ccnd1* (cyclin D1), *Ctgf* (Connective tissue growth factor), *Gpc3* (Glypican-3), and *Igf2* (Insulin-like growth factor-2); the onco-foetal markers *Afp* and *H19*; and the cancer stem cell markers *Klf4* (Kruppel-like factor 4), *Aldh1* (Aldehyde dehydrogenase 1), and *Cd133* (CD133/Prominin-1) (**Figure 4D**). This suggested that young *Otulin*^Δhep^ mice could be prone to development of HCC.

**Figure 4.**
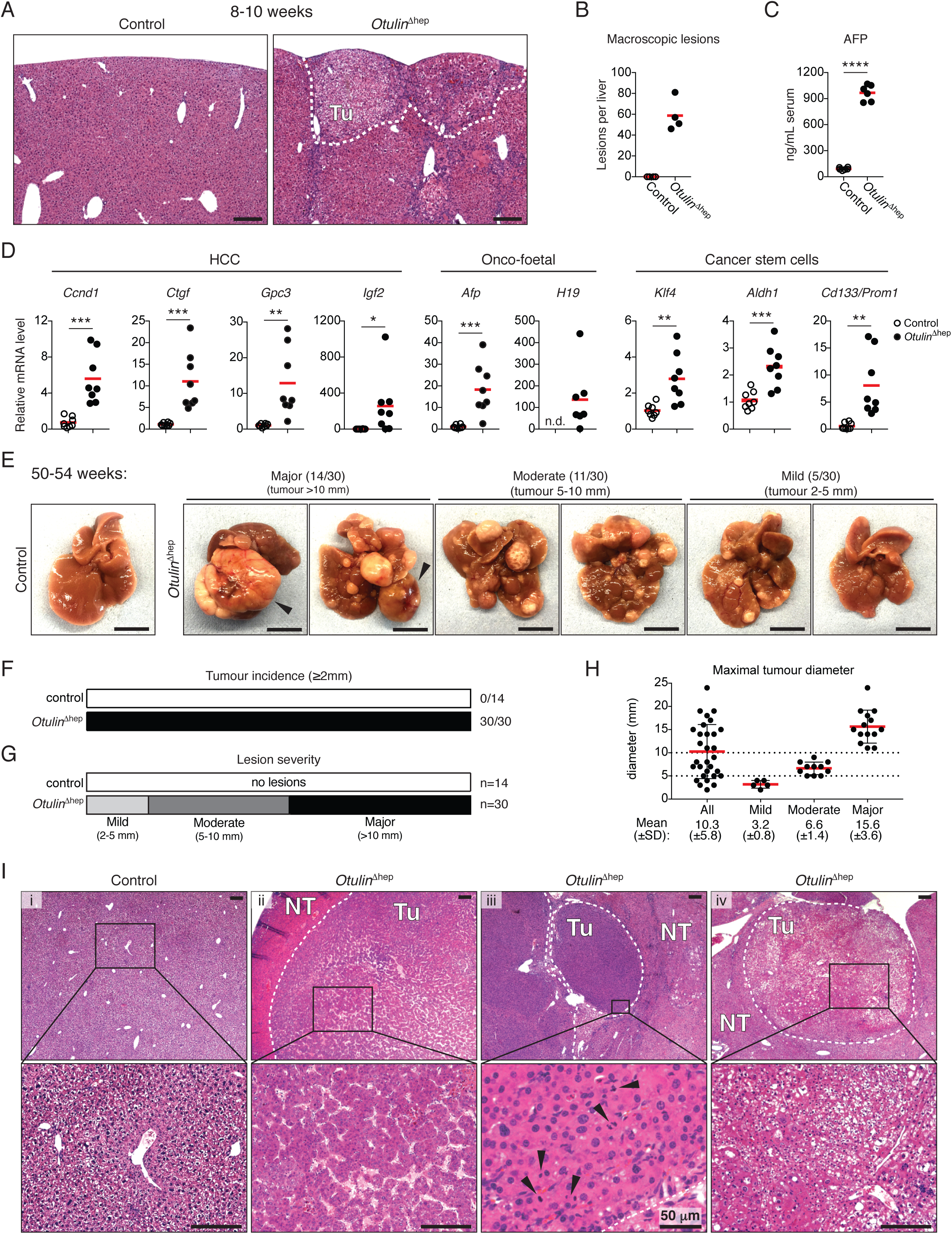
Hepatocellular carcinoma in *Otulin*^Δhep^ mice. **(A)** Micrographs of H&E stained liver sections from *Otulin*^Δhep^ and control mice aged 8-10 weeks. The dotted line indicates two subcapsular tumours. Micrographs are representative of eight mice of each genotype. Tu, tumour. **(B)** Quantification of the number of macroscopically discernible lesions (tumours, nodules, and discolourations) in *Otulin*^Δhep^ and control mice aged 8-10 weeks. Data are representative of four mice of each genotype. **(C)** Analysis of AFP levels in serum from terminal bleeds of *Otulin*^Δhep^ (n=6) and control (n=6) mice aged 8-10 weeks. **(D)** Relative mRNA expression of the indicated cancer markers in livers from *Otulin*^Δhep^ (n=8) and control (n=8) aged 8-10 weeks measured by quantitative RT-PCR. **(E)** Representative macroscopic appearance of *Otulin*^Δhep^ and control livers at the age of 50-54 weeks, grouped by severity. Arrowheads indicate highly vascularised tumours. Scale bars indicate 1 cm. **(F-G)** Quantification of the number of mice with the presence of a tumour ≥2 mm in diameter (F) or the number of mice in each severity group as indicated (G) in *Otulin*^Δhep^ and control mice at the age of 50-54 weeks. **(H)** Maximal tumour size in *Otulin*^Δhep^ mice, grouped by degree of pathology. Each data point represents the maximal tumour size in one mouse. Red bars indicate means ±SD. **(I)** Micrographs of H&E stained liver sections from *Otulin*^Δhep^ and control mice aged 50-54 weeks. (i) shows control livers, (ii) shows HCC with abnormal macrotrabecular pattern, (iii) shows raised nuclear-cytoplasmic ratio, irregular nuclear outlines and several mitotic figures (arrowheads), and (iv) shows a tumour with a steatohepatic appearance, with also enlarged and irregular nuclei. Micrographs are representative of four controls and 15 *Otulin*^Δhep^ mice. Tu, tumour. NT, non-tumour. **(B-D)** Data are presented as individual data points, each representing one mouse. Red bars indicate means. Data were analysed using an unpaired, two-sided Student’s *t* test. n.s., non-significant. See also Figure S4.

Indeed, dissection of livers from *Otulin*^Δhep^ mice aged 50-54 weeks revealed the presence of multiple large tumours (**Figure 4E**). The tumour incidence (presence of a tumour ≥2 mm) was 100% in *Otulin*^Δhep^ mice while no lesions were observed in controls (**Figure 4F**). The tumour size, number, and severity were variable in *Otulin*^Δhep^ mice. Nearly half of them presented with major pathology (presence of tumour >10 mm in diameter; 14/30) (**Figure 4G)**, many of which had highly vascularised tumours (**Figures 4E, arrowheads, and S4A**). Approximately one third of the *Otulin*^Δhep^ mice developed moderate pathology (presence of tumour 5-10 mm in diameter; 11/30), while only a few mice developed mild pathology (presence of tumour 2-5 mm in diameter; 5/30) (**Figures 4E and 4G**). The average maximal tumour size in the *Otulin*^Δhep^ livers was 10.3 mm (±5.3 mm), 3.2 mm (±0.8 mm) for mice with mild pathology, 6.6 mm (±1.4 mm) for moderate, and 15.6 mm (±3.6 mm) for mice with major pathology (**Figure 4H**). Microscopic examination uncovered the presence of malignant tumours corresponding to well and moderately differentiated HCC (**Figure 4I**) (Wanless and Party, 1995). The analysed tumours were characterised by expansive growth and the absence of portal tracts (**Figure 4I**), broad trabecular growth (>4 cells wide) (**Figure 4I**, column ii), increased eosinophilia (columns ii and iii) or cell clearance (column iv), increased number of mitotic figures (column iii, arrowheads), as well as high pleomorphism and atypical nuclei (column iv), all indicative of malignant HCC (Wanless and Party, 1995). Occasionally, tumours also showed focal necrosis and cystic degeneration (**Figure S4B**), indicating fast-growing and aggressive tumours. Pre-malignant dysplatic nodules with severe anisokaryosis and atypic nuclei were also present (Kojiro and Roskams, 2005). Analysis of *Otulin*^Δhep^ mice aged 32 weeks revealed moderate pathology (**Figure S4C**) and the presence of well differentiated tumours (**Figure S4D**), occasionally with indications of poor demarcation and the absence of portal tracts, indicating that these are early neoplastic tumours. This suggests that malignancy arises between 32 and 50 weeks of age in *Otulin*^Δhep^ mice.

Collectively, these data show that deletion of *Otulin* in hepatocytes leads to spontaneous development of hepatocellular carcinoma, demonstrating that OTULIN acts as a tumour suppressor in the liver.

### Steatohepatitis in *Otulin*^Δhep^ mice is independent of TNFR1 signalling

TNF is the key driver of inflammation in both ORAS patients and ORAS mouse models with OTULIN-deficiency in immune cells (Damgaard et al., 2016; 2019; Zhou et al., 2016). Dysregulated TNFR1 signalling also contributes to development of liver disease and cancer (Luedde et al., 2014), and liver-specific deletion of CYLD, the other M1-polyUb-regulating DUB, causes TNFR1-mediated hepatitis, fibrosis, and HCC (Nikolaou et al., 2012). We therefore investigated if TNFR1 signalling contributes to development of the NASH-like disease in *Otulin*^Δhep^ mice. Surprisingly, co-deletion of *Tnfr1* (p55-TNFR1) in *Otulin*^Δhep^ mice did not prevent the development of liver disease (**Figures 5A and S5A**). *Otulin*^Δhep^ and *Otulin*^Δhep^;*Tnfr1^−/−^* mice aged 8-12 weeks developed indistinguishable pathology with similar numbers of macroscopic lesions (**Figures 5A-C**). Microscopic examination revealed virtually identical abnormal histology with dysplastic nodules, large cell change, anisokaryosis, and cytoplasmic inclusions in both *Otulin*^Δhep^ and *Otulin*^Δhep^;*Tnfr1^−/−^* mice (**Figures 5C, top panels, and S5B**). The extent of collagen deposition and the pattern of bridging fibrosis was also unchanged by the deletion of TNFR1 (**Figures 5C, bottom panels, and 5D**). Serum levels of ALT and AST, which reflect the degree of cell death in the liver parenchyma (Luedde et al., 2014), were not significantly altered in the *Otulin*^Δhep^;*Tnfr1^−/−^* mice either (**Figure 5E**). We therefore conclude that the cellular aberrations leading to NASH-like liver disease in young adult *Otulin*^Δhep^ mice are independent of signalling through TNFR1 and therefore distinct from the pathology in CYLD-deficient livers (Nikolaou et al., 2012).

**Figure 5.**
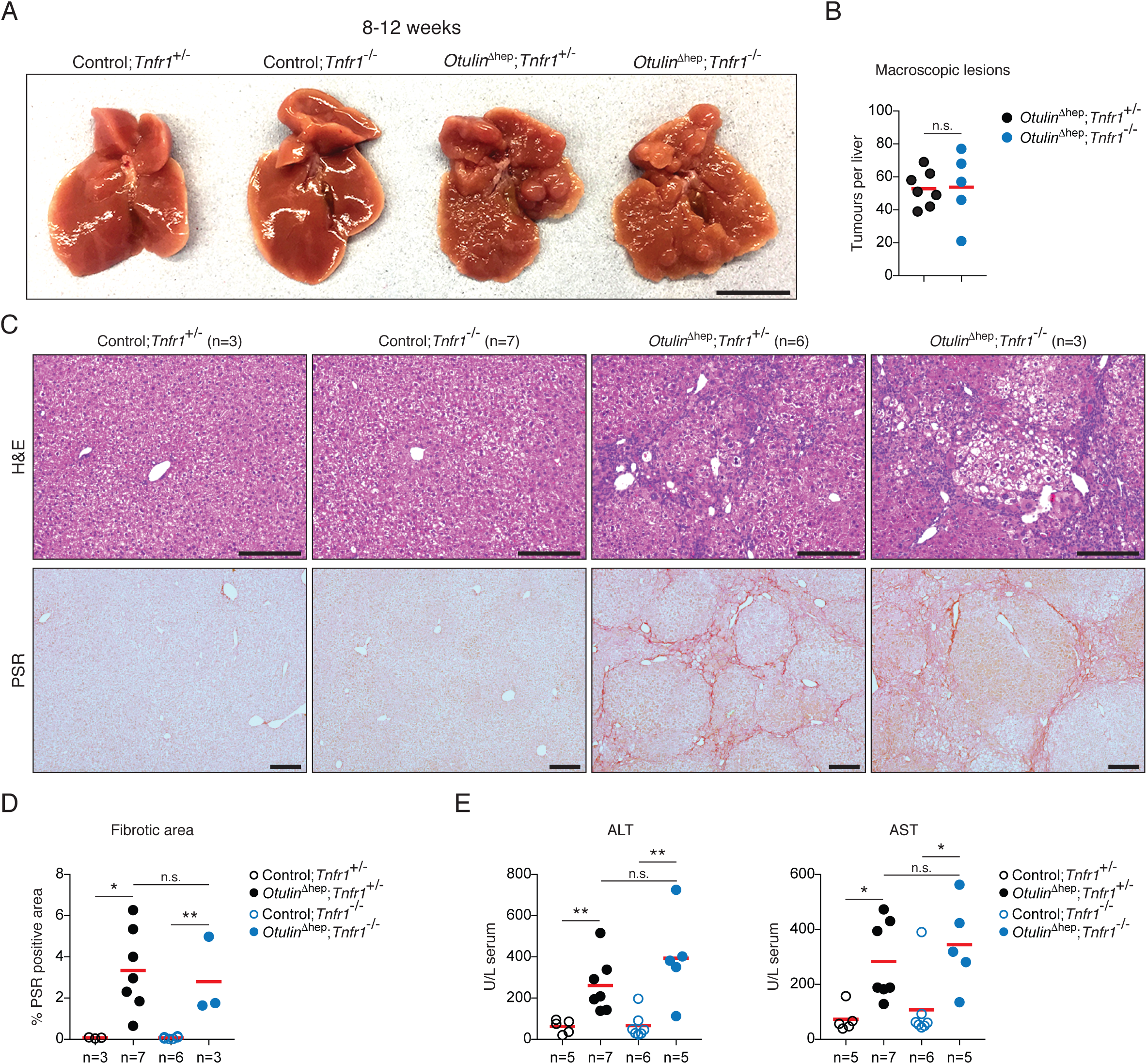
Liver disease in *Otulin*^Δhep^ mice is independent of TNFR1 signalling. **(A)** Representative macroscopic appearance of livers from *Otulin*^Δhep^ mice, *Otulin*^Δhep^;*Tnfr1^−/−^* mice, and their respective controls at the age of 8-12 weeks. Scale bar indicates 1 cm. **(B)** Quantification of the number of macroscopically discernible lesions (tumours, nodules, and discolourations) in *Otulin*^Δhep^ (n=7) and *Otulin*^Δhep^;*Tnfr1^−/−^* (n=5) mice aged 8-12 weeks. **(C)** Micrographs of liver sections from *Otulin*^Δhep^ mice, *Otulin*^Δhep^;*Tnfr1^−/−^* mice, and their respective controls at the age of 8-12 weeks stained with H&E or PSR as indicated. **(D)** Quantification of PSR-positive (fibrotic) area in liver sections *Otulin*^Δhep^ mice, *Otulin*^Δhep^;*Tnfr1^−/−^* mice, and their respective controls at the age of 8-12 weeks. **(E)** Analysis of ALT and AST levels in serum from terminal bleeds of *Otulin*^Δhep^ mice, *Otulin*^Δhep^;*Tnfr1^−/−^* mice, and their respective controls at the age of 8-12 weeks. **(B, D-E)** Data are presented as individual data points, each representing one mouse. Red bars indicate means. Data were analysed using unpaired, two-sided Student’s *t* tests. n.s., non-significant. See also Figure S5.

### Prominent neonatal steatosis and aberrant mTOR activation in *Otulin*^Δhep^ mice

As the NASH-like phenotype in young adult *Otulin*^Δhep^ mice was independent of TNFR1 signalling, we examined livers from younger *Otulin*^Δhep^ mice to explore the onset of the phenotype in these mice and how it presented. Analysis of livers from neonatal *Otulin*^Δhep^ and control mice at postnatal day (P) 3 and P9 showed that OTULIN ablation was efficient at this age and that HOIP expression was reduced (**Figures S6A-D**), similar to our observations at 8-10 weeks. Strikingly, we found that *Otulin*^Δhep^ mice displayed marked steatosis at P3 and P9 (**Figures 6A-B and S6E**). The neonatal *Otulin*^Δhep^ livers were pale and appeared oily, particularly at P9 (**Figure 6A**). The level of total cholesterol in serum was also increased at P9, while triglyceride and glucose levels were similar between *Otulin*^Δhep^ and control mice (**Figure 6C**). Histopathological examination revealed progressive lipid accumulation, mainly microsteatosis, between P3 and P9 in *Otulin*^Δhep^ mice (**Figures 6B, top and centre panels, and S6E**). Lipid-specific Oil Red O staining confirmed prominent steatosis and showed a significant increase in both the total amount of fat and the number of lipid droplets present in the P9 livers (**Figures 6B, bottom panels, and 6D**). The progression of the *Otulin*^Δhep^ phenotype, from steatosis to steatohepatitis, fibrosis, and HCC, therefore closely resembles that of patients with non-alcoholic fatty liver disease (NAFLD) and NASH (Farrell and Larter, 2006; Llovet et al., 2016).

**Figure 6.**
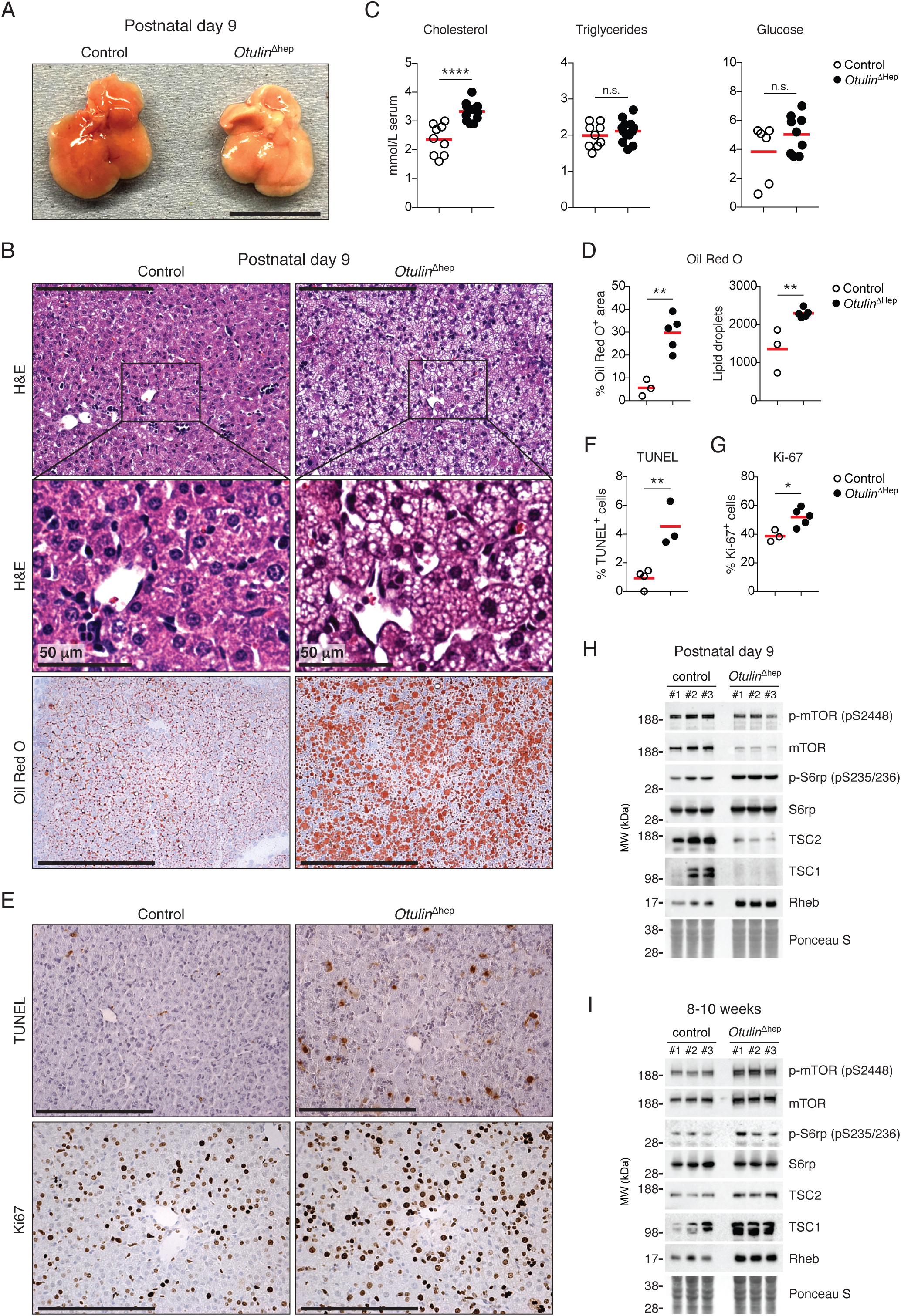
Steatosis and aberrant mTOR activation in neonatal *Otulin*^Δhep^ mice. **(A)** Representative macroscopic appearance *Otulin*^Δhep^ and control livers at the age of 9 days. Scale bar indicates 1 cm. **(B)** Micrographs of liver sections from *Otulin*^Δhep^ and control mice at the age of 9 days stained with H&E and Oil Red O as indicated. H&E staining shows pale hepatocytes with varying sized vacuoles in *Otulin*^Δhep^ mice, which is confirmed as fat by Oil Red O stainig. Micrographs are representative of seven controls and six *Otulin*^Δhep^ mice for H&E, and three controls and five *Otulin*^Δhep^ mice for Oil Red O. **(C)** Analysis of cholesterol, triglyceride, and glucose levels in serum from terminal bleeds of *Otulin*^Δhep^ (n=9) and control (n=6) mice at the age of 9 days. **(D)** Quantification of Oil Red O-positive area (left) and number of lipid droplets (right) in liver sections *Otulin*^Δhep^ (n=5) and control (n=3) at the age of 9 days as shown in (B). **(E)** TUNEL (top panels) and anti-Ki67 (bottom panels) stainings of liver sections from *Otulin*^Δhep^ and control mice aged 9 days. Data are representative of four control and three *Otulin*^Δhep^ mice for TUNEL, and three controls and five *Otulin*^Δhep^ mice for Ki67. **(F-G)** Quantification of TUNEL-(F) and Ki67-positive (G) cells in liver from *Otulin*^Δhep^ and control at the age of 9 days as shown in (E). TUNEL (F), *Otulin*^Δhep^ (n=3) and control (n=4), and anti-Ki67 (G), *Otulin*^Δhep^ (n=5) and control (n=3). **(H-I)** Immunoblot analysis of mTOR pathway components and activation in whole-liver lysate from three *Otulin*^Δhep^ mice and three controls aged 9 days (H) and 8-10 weeks (I). **(C-D, F-G)** Data are presented as individual data points, each representing one mouse. Red bars indicate means. Data were analysed using an unpaired, two-sided Student’s *t* test. n.s., non-significant. See also Figure S6.

Immunohistochemical analysis showed that the proportion of TUNEL-positive dead cells in the *Otulin*^Δhep^ livers was significantly increased at P9 (**Figures 6E, top panels, and 6F**), comparable to the increase observed at 8-10 weeks. Both at P3 and P9, caspase-3 cleavage and activation was increased (**Figures S6A and S6C**), supporting our previous observations of apoptotic cell death in OTULIN-deficient livers. The number of Ki67-positive proliferating cells also increased in the P9 *Otulin*^Δhep^ livers, although only marginally (**Figures 6E, bottom panels, and 6G**), likely due to the fact that the liver at this age is a highly proliferative organ already. In contrast, we did not detect any signs of collagen deposition and fibrosis at either P3 or P9 in these mice (**Figures S6F and S6G**).

The kinase mTOR is a master regulator of cellular metabolism and growth (Mossmann et al., 2018), and increased mTOR activity promotes liver carcinogenesis in mice (Guri et al., 2017; Menon et al., 2012). In models of mTOR-driven carcinogenesis, metabolic alterations accompany hepatocyte damage and proliferation (Guri et al., 2017; Menon et al., 2012), akin to our observations in *Otulin*^Δhep^ mice. This prompted us to investigate if mTOR signalling was altered in *Otulin*^Δhep^ livers. Intriguingly, we found that mTOR signalling was increased in P9 *Otulin*^Δhep^ livers compared with controls (**Figures 6H and S6D**). When compared with the overall lower expression of total mTOR protein in liver lysates from P9 *Otulin*^Δhep^ mice, the relative phosphorylation of the activating Ser2448 of mTOR was increased compared with controls (**Figure 6H**). This correlated with increased phosphorylation of the mTOR-dependent substrate S6 ribosomal protein (S6rp) (**Figure 6H**), which was also detected in P3 livers (**Figure S6H**). Phosphorylation of mTOR and S6rp was associated with reduced levels of the TSC complex (consisting of TSC1 and TSC2), a negative regulator of mTOR, and an increased expression of the mTOR activator Rheb in OTULIN-deficient livers (**Figures 6H and S6H**). At the age of 8-10 weeks, *Otulin*^Δhep^ livers showed more normal, but still slightly increased mTOR activation (**Figures 6I and S6I**). The expression of the TSC complex was comparable to controls, but Rheb expression was still increased (**Figure 6I**). This correlated with a slight increase in mTOR and S6rp phosphorylation (**Figure 6I**), albeit to a lesser extent than observed at P3 and P9. OTULIN deficiency in hepatocytes therefore leads to altered expression of components of the mTOR pathway and increased mTOR activation, revealing a potential role for OTULIN and M1-polyUb in regulation of mTOR signalling.

### mTOR inhibition reduces liver disease in *Otulin*^Δhep^ mice

In humans, mTOR signalling is upregulated in 40-50% of HCC cases and is associated with poor prognosis (Matter et al., 2014). Multiple mTOR inhibitors, mostly derivatives of the allosteric inhibitor rapamycin, are therefore being trialled for treatment of HCC (Matter et al., 2014). In mice, increased mTOR signalling in TSC1-deficient or TSC1-PTEN-double-deficient livers leads to HCC development, and the liver pathology and tumorigenesis in these genetic models of mTOR-driven HCC can be reduced by treatment with mTOR inhibitors (Guri et al., 2017; Menon et al., 2012). To examine if OTULIN deficiency led to mTOR-driven liver disease, we tested if pharmacological inhibition of mTOR could reduce the pathology observed in the *Otulin*^Δhep^ mice. As altered mTOR activation is already evident together with steatosis at P3, we treated *Otulin*^Δhep^ mice with rapamycin from birth until the age of 8 weeks. Rapamycin treatment was not well tolerated in *Otulin*^Δhep^ mice. Treated *Otulin*^Δhep^ mice displayed markedly reduced weight gain when compared with vehicle-treated mice and even rapamycin-treated controls (**Figure S7A**), demonstrating a pharmacogenetic interaction between OTULIN deficiency and mTOR inhibition. The condition of the rapamycin-treated *Otulin*^Δhep^ mice meant that for many mice the experiment had to be stopped at 6 weeks of age (**Figure S7A**).

Remarkably, despite early termination of the experiment, rapamycin treatment caused an improvement of liver pathology in *Otulin*^Δhep^ mice compared with vehicle-treated *Otulin*^Δhep^ mice of the same age (**Figure 7A**). Rapamycin treatment reduced both the number and size of macroscopic lesions in the livers of *Otulin*^Δhep^ mice, but did not completely prevent liver disease (**Figures 7A-C**). The livers from the rapamycin-treated *Otulin*^Δhep^ mice appeared smaller than vehicle-treated *Otulin*^Δhep^ mice or rapamycin-treated controls (**Figure 7A**), but relative to body weight they were not different from vehicle-treated *Otulin*^Δhep^ livers (**Figure S7B**). Microscopically, rapamycin reduced the histological abnormalities and the number of dysplastic foci and nodular growths in the *Otulin*^Δhep^ livers (**Figures 7C and S7C**). Hepatocyte dysplasia and the inflammatory cells in the parenchyma were decreased in the rapamycin-treated *Otulin*^Δhep^ livers, although some of the cellular changes, including atypical nuclei and hepatocyte hypertrophy, persisted (**Figure 7C, inserts**). In addition, rapamycin treatment significantly reduced the collagen deposition and fibrosis in *Otulin*^Δhep^ mice (**Figures 7C-C**). Importantly, ALT and AST levels in serum were not reduced by mTOR inhibition (**Figure 7E**), suggesting that apoptosis in *Otulin*^Δhep^ livers is independent of aberrant mTOR activation. Our findings demonstrate that mTOR activity promotes tumourigenesis and fibrosis in *Otulin*^Δhep^ mice, but also that mTOR inhibition by rapamycin is insufficient to completely prevent liver pathology in these mice.

**Figure 7.**
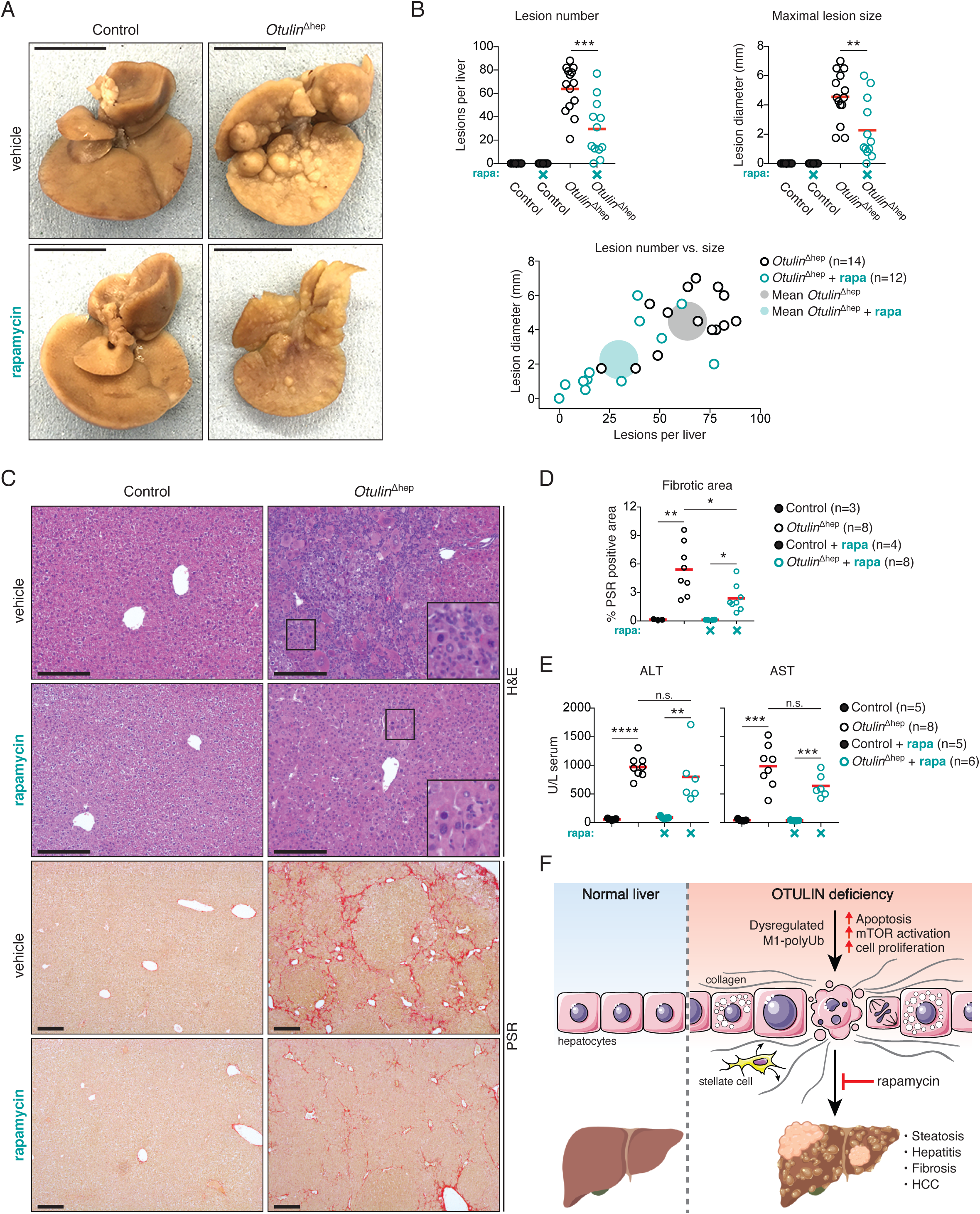
The mTOR inhibitor rapamycin reduces liver pathology in *Otulin*^Δhep^ mice. **(A)** Representative macroscopic appearance of formalin-fixed livers from *Otulin*^Δhep^ and control mice at the age of six weeks treated with rapamycin or vehicle as indicated. Scale bar indicates 1 cm. **(B)** Quantification of and correlation between the number and maximal size of macroscopically discernible lesions (tumours, nodules, and discolourations) in *Otulin*^Δhep^ and control mice aged 6-8 weeks treated with rapamycin (rapa) or vehicle as indicated. Data were pooled from two independent experiments. Opaque circles indicate the mean lesion number and maximal lesion size of the two groups. Red bars indicate means. **(C)** Micrographs of liver sections from *Otulin*^Δhep^ and control mice at the age of six weeks treated with rapamycin or vehicle and stained with H&E and PSR as indicated. Data are representative of three vehicle-treated controls, eight vehicle-treated *Otulin*^Δhep^ mice, four rapamycin-treated controls, and eight rapamycin-treated *Otulin*^Δhep^ mice. Inserts show cellular changes at high magnification. **(D)** Quantification of PSR-positive (fibrotic) area in liver sections from *Otulin*^Δhep^ and control mice at the age of six weeks treated with rapamycin (rapa) or vehicle as indicated. Red bars indicate means. **(E)** Analysis of ALT and AST levels in serum from terminal bleeds from control and *Otulin*^Δhep^ mice at the age of 6-8 weeks treated with vehicle or rapamycin (rapa) as indicated. Data were pooled from two independent experiments. **(F)** Model of the cellular and phenotypic changes in OTULIN-deficient livers. **(B,D-E)** Data are presented as individual data points, each representing one mouse. Data were analysed using unpaired, two-sided Student’s *t* tests. n.s., non-significant. See also Figure S7.

## Discussion

We provide genetic evidence that OTULIN is a tumour suppressor and crucial *in vivo* regulator of liver homeostasis in mice, identify mTOR signalling as a surprising driver of liver disease in OTULIN-deficient mice, and show that mTOR inhibition with rapamycin can improve liver pathology caused by OTULIN deficiency.

OTULIN mutations in humans cause a severe autoinflammatory syndrome, ORAS (Damgaard et al., 2016; 2019; Nabavi et al., 2019; Zhou et al., 2016), and genetic ablation of OTULIN in immune cells in mice replicate many inflammatory hallmarks of ORAS (Damgaard et al., 2016). The discovery that OTULIN is a tumour suppressor and vital for maintaining liver homeostasis highlights just how critically important proper regulation of M1-polyUb signalling is and expands the range of pathologies associated with OTULIN deficiency. Inflammatory disease in ORAS patients is successfully ameliorated by TNF-blocking agents, such as Infliximab or Etanercept (Damgaard et al., 2016; 2019). However, co-deletion of TNFR1 has no bearing on the liver pathology in *Otulin*^Δhep^ mice. This clearly indicates that the liver pathogenesis is independent of TNF signalling and suggests that liver function should be monitored closely in ORAS patients, both for development of spontaneous disease and for hypersensitivity to pharmaceuticals or food-borne hazards.

Phenotypic and molecular analyses of the OTULIN-deficient livers revealed early-onset progressive liver pathology with temporarily distinct phases in *Otulin*^Δhep^ mice. Within days of birth, *Otulin*^Δhep^ mice develop steatosis, which by the age of eight weeks has developed into steatohepatitis with severe fibrotic lesions and pre-malignant tumours, and HCC develops within 7-12 months in the *Otulin*^Δhep^ livers. This phenotypic appearance and pattern of disease progression is remarkably similar to the advancement of liver disease in human NASH patients. Similar to NASH in humans, pathology in *Otulin*^Δhep^ mice is also associated with apoptosis and compensatory hepatocyte proliferation (**Figure 7F**) (Farrell and Larter, 2006; Luedde et al., 2014). LUBAC levels in *Otulin*^Δhep^ livers are reduced, similar to the effect of OTULIN deficiency in fibroblasts, which are sensitised to TNF-induced apoptosis (Damgaard et al., 2019). Apoptosis in *Otulin*^Δhep^ livers is evident as caspase-3 activation from P3 and through to development of steatohepatitis and fibrosis at eight weeks, suggesting that apoptosis is a driver of liver pathology in these livers, similar to CYLD- or NEMO-deficient livers (Luedde et al., 2007; Nikolaou et al., 2012). Surprisingly, unlike CYLD-deficient livers (Nikolaou et al., 2012), liver damage in *Otulin*^Δhep^ livers is independent of TNFR1 signalling, suggesting that the liver diseases caused by deficiency in the two main M1-polyUb-regulating DUBs have distinct pathogeneses. This raises the question of what the inducer of apoptosis in OTULIN-deficient hepatocytes is. Delineation of the pathway(s) to TNFR1-independent apoptosis in OTULIN-deficient livers will be important to understand the pathogenesis of steatohepatitis and HCC development in *Otulin*^Δhep^ mice.

*Otulin*^Δhep^ mice appear healthy until at least 12 months of age. However, they exhibit obvious liver pathology from shortly after birth. The degree of disease in OTULIN-deficient livers, both in young mice with NASH and older mice with HCC, is unusually severe compared with other regulators of M1-polyUb signalling. Liver-specific deletion of A20, which binds and regulates M1-polyUb signalling via its ZnF7 domain (Verhelst et al., 2012), causes low grade inflammation, but no fibrosis or spontaneous HCC development (Catrysse et al., 2016). More similar to OTULIN-deficiency, CYLD inactivation in liver causes progressive fibrosis and HCC by 12 months of age, but the macroscopic appearance of CYLD-deficient livers, the pattern of collagen deposition, and dependency on TNFR1 signalling is markedly different from *Otulin*^Δhep^ mice (Nikolaou et al., 2012). Deletion of HOIP, the catalytic subunit of LUBAC (Kirisako et al., 2006), also causes liver inflammation. Intriguingly, HOIP-deficient livers are not reported to display signs of neonatal steatosis or fibrosis, and only a fraction of livers show severe pathology and HCC by 18 months (Shimizu et al., 2017). Even though OTULIN deficiency – or lack of its catalytic activity – may phenocopy HOIP deficiency in some respects (Heger et al., 2018; Peltzer and Walczak, 2019), our finding of early-onset, highly penetrant, and severe liver disease in *Otulin*^Δhep^ mice implies important pathophysiological differences between complete loss of M1-linked polyUb signalling in HOIP-deficient livers and dysregulated M1-polyUb signalling in OTULIN-deficient livers.

Our examination of neonatal *Otulin*^Δhep^ mice revealed an unexpected phenotype of steatosis, hypercholesterolaemia, and increased mTOR signalling in OTULIN-deficient livers. To our knowledge, this is the first report of a link between M1-polyUb and mTOR signalling. OTULIN-deficiency appears to cause changes in the expression of mTOR regulators such as the TSC complex and Rheb in the liver, likely leading to aberrant mTOR activation. However, the molecular nature of this dysregulation, and whether it is a direct or indirect effect, is unclear. More mechanistic studies are needed to elucidate how OTULIN and M1-polyUb may regulate mTOR complexes and signalling. Intriguingly, by eight weeks of age, lipid and glucose levels in serum are reduced. These reductions are accompanied by a striking decrease in liver glycogen content, indicating that OTULIN deficiency may cause defects in carbohydrate storage or metabolism that could fuel liver disease. The reduced glycogen storage and altered metabolism may also provide an explanation for the reduced tolerance to rapamycin treatment in *Otulin*^Δhep^ mice, despite the improvement of overall liver pathology.

Collectively, our findings demonstrate that OTULIN is a tumour suppressor that prevents cell death, inflammation, and metabolic derangements in liver (**Figure 7F**), highlighting how delicately balanced M1-polyUb signalling needs to be to prevent disease.

## Acknowledgements

We would like to thank Irina Pshenichnaya (Cambridge Stem Cell Institute) for technical advice, Prof. Kazuhiro Iwai (Kyoto University) for sharing the HOIP antibody, the MRC LMB Genotyping Service for mouse genotyping, and Shona Butler, Graham “Ledge” Ledger, Richard Berks, Jennifer Roe, and MRC ARES and Biomed staff for assistance with animal experiments. Work in the D.K. laboratory was supported by the Medical Research Council [U105192732], the European Research Council [724804], and the Lister Institute for Preventive Medicine. R.B.D was supported by a Marie Sklodowska Curie Individual Fellowship from the European Commission [MC-IF-654019], a Lundbeckfonden Postdoctoral Fellowship [R232-2016-1904/R265-2017-2998], and a Research Fellowship from Corpus Christi College Cambridge. A.N.J.M. was supported by the Medical Research Council [U105178805]. Figure 7F was produced using Servier Medical Art and Biorender.

## Author contributions

R.B.D. conceived the project. R.B.D., A.N.J.M., and D.K. coordinated the research. R.B.D. designed and performed the experiments, assisted by H.E.J. R.B.D. analysed and interpreted the data, assisted by M.E.D.A and S.E.D. R.B.D. and D.K. wrote the paper with input from all authors.

## STAR Methods

### CONTACT AND MATERIALS AVAILABILITY

Further information and requests for resources and reagents should be directed to and will be fulfilled by the Lead Contact, David Komander.

### EXPERIMENTAL MODEL AND SUBJECT DETAILS

#### Mice

All animal experiments were undertaken with the approval of the United Kingdom Home Office and the MRC Centre Ethical Review Committee and performed in accordance with the Animals (Scientific Procedures) Act of 1986. All mice were on a C57BL/6 or B6.SJL background and maintained under specific pathogen-free conditions in individually ventilated cages (Techniplast GM500, Techniplast) on Lignocel FS14 spruce bedding (IPS, Ltd.) with environmental enrichment (fun tunnel, chew stick, and Enviro-Dri® nesting material (LBS)) at 19-23 °C with light from 7.00 a.m. to 7.00 p.m. and fed Dietex CRM pellets (Special Diet Services) ad libitum.

Mice carrying the *loxP*-flanked (flox) or deleted (del) *Otulin* alleles and the *Otulin*-*Rosa26*-Cre-ERT2 inducible knockout strain were described previously (Damgaard et al., 2016). The *Otulin*^Δhep^ strain with conditional deletion of OTULIN in hepatocytes was generated by breeding *Otulin*^flox/flox^ and *Otulin*^del/flox^ mice with mice expressing the Cre recombinase under the control of a serum albumin promoter (*Alb*-Cre) (Postic et al., 1999) obtained from The Jackson Laboratory (Bar Harbor, ME). *Otulin*^Δhep^ mice used in this study were either *Otulin*^flox/flox^;*Alb*-Cre^Tg+^ or *Otulin*^del/flox^;*Alb*-Cre^Tg+^. Control mice were mostly *Otulin*^+/flox^;*Alb*-Cre^Tg+^ or occasionally wild type C57BL/6. *Otulin*^Δhep^ mice were bred with *Tnfrsf1a*^tm1Imx^ mice (Peschon et al., 1998) (Jackson Laboratory, Bar Harbor, ME) to generate the *Otulin^Δ^*^hep^;*Tnfr1*^−/−^ double-deficient mice. All mice were genotyped by PCR.

In individual experiments, mice were matched for age and sex whenever possible. For *Otulin*-*Rosa26*-Cre-ERT2 bone marrow chimeric mice, recipient mice were 4-5 months old and in each experiment were either all male or all female. For *Otulin*^Δhep^ mice and corresponding controls, experiments were performed with a mix of male and female mice, age-matched between three days and 54 weeks as indicated in figures and legends. Mice were allocated to experimental groups based on genotype. Sample sizes were estimated using pilot experiments. All mice were culled by exposure to rising levels of CO2 followed by exsanguination.

### METHOD DETAILS

#### Generation of bone marrow chimeric mice

From one week before transplantation until two weeks after the transplantation, the drinking water was supplemented with 0.1 mg/mL enrofloxacin (Baytril®; Bayer). Bone marrow cells (2 × 10^6^) from wild type CD45.1^+^ B6/SJL mice, isolated by flushing femurs with sterile PBS supplemented with 2% foetal bovine serum (FBS; Gibco), were resuspended in 100 μl PBS and then injected intravenously (i.v.) into 4-5 months old sex- and age-matched *γ*-irradiated *Rosa26-*Cre-ERT2-*Otulin*^+/flox^ or -*Otulin*^del/flox^ recipients (given two doses of 4.5 Gy). Six weeks after reconstitution, mice were given three doses of tamoxifen (Sigma-Aldrich, St Louis, MO; 1 mg in sunflower oil with 10% ethanol per dose) i.p. over three days. Mice were closely monitored and weighed daily. At 8-10 days after the first tamoxifen dose, mice were culled and samples taken for analyses.

#### Rapamycin treatment of *Otulin*^Δhep^ mice

*Otulin*^Δhep^ and control (*Otulin*^+/flox^;*Alb*-Cre^Tg+^) mice were bred by timed matings. Dams pregnant with pups to be allocated to rapamycin-treated groups received one intraperitoneal (i.p.) injection of rapamycin (1 mg/kg) at E17.5. After birth, pups were allocated to experimental groups based on their genotype and fostered onto pseudopregnant CD-1 mothers. At postnatal day 3 (P3), lactating CD-1 foster mothers received one subcutaneous (s.c.) dose of rapamycin (1 mg/kg) or vehicle. From P8, pups were injected i.p. with rapamycin or vehicle twice weekly until 8 weeks of age, or until a humane endpoint. Mice received increasing doses of rapamycin as follows: postnatal day 3 (P3), 20 μg; P11, 25 μg; P15, 30 μg; P18, 35 μg; P22, 135 μg; P25, 180 μg; P29, 240 μg; P32, 240 μg; P36, 300 μg; P39, 300 μg; P43, 330 μg; P46, 330 μg; P50, 330 μg; P53 360 μg; P57, 360 μg; equivalent to 1 mg/kg between P8 and P18 and 3 mg/kg from P22 until the end of the experiment based on the predicted weight of the mice at the time of each injection. Mice that met a humane endpoint before the age of 39 days were excluded from analysis. Rapamycin (LC Laboratories, Woburn, MA) was dissolved in 70% ethanol at 20 mg/mL and diluted to 0.2-0.6 mg/mL in sterile PBS containing 0.5% (v/v) Tween-80 (VWR, Lutterworth, UK) and 0.5% (v/v) PEG-400 (Hampton Research, Aliso Viejo, CA) before injection.

#### Blood cell counts

Whole blood from terminal bleeds was collected in EDTA-containing Blood Collection Tubes (Greiner) and analysed on a scil Vet abcPlus^+^ haematological analyser (scil Animal Care Company, Gurnee, IL).

#### Histology

Tissue samples were dissected and placed in ice-cold PBS containing 3% (v/v) FCS. For fresh frozen sections, samples were embedded in OCT Embedding Medium (Thermo Scientific), left 2-3 min to acclimate to the OCT, and then frozen by submerging the cryomould in dry ice-cooled isopropanol before transferring the moulds to −80 °C for storage before sectioning. For sections of fixed tissue, samples were transferred to 10% neutral buffered formalin (Sigma) and fixed for 24 h at room temperature. Fixed tissues were paraffin embedded, sectioned (5 μm), and stained at AML Laboratories, Inc., Jacksonville, FL, or Cambridge Stem Cell Institute Histology Core Facility, University of Cambridge, Cambridge, UK, according to standard procedures. Haematoxylin and Eosin (H&E) staining: after deparaffinisation of formalin-fixed paraffin-embedded (FFPE) sections with xylene and rehydration, slides were stained with Harris Haematoxylin for 3 min, differentiated with acidified ethanol, and post-stained with Eosin for 30 sec, followed by dehydration and clearance. Slides were then mounted with DPX mounting medium. Picro Sirius Red (PSR) staining: after deparaffinisation of FFPE sections with xylene and rehydration, slides were stained with a freshly prepared Weigert’s Haematoxylin solution for 4-8 min followed by staining with Picro Sirius Red (Sirius red F3B) (Sigma) dissolved in a saturated aqueous solution of picric acid for 1 h. Slides were then rinsed in acidified water, dehydrated, cleared, and mounted with DPX mounting medium. Periodic acid–Schiff (PAS) stain: after deparaffinisation of FFPE sections with xylene and rehydration, slides were oxidised in 0.5% periodic acid solution for 5 min, rinsed in distilled water, and stained with Schiff reagent for 15 min. Slides were then washed, counterstained with Haematoxylin, dehydrated, cleared, and mounted with DPX mounting medium. Oil Red O staining: fresh-frozen unfixed sections were stained with Oil Red O (Sigma) dissolved in 2-propanol for 10 min, quickly rinsed in tap water, and followed by counterstaining with Mayer’s Hematoxylin and prolonged wash in the tap water. Slides are then mounted with glycerol and imaged immediately.

#### Immunohistochemistry and TUNEL assay

Immunohistochemistry (IHC) was performed on deparaffinised and rehydrated FFPE sections. Antigen retrieval was performed in citric acid buffer, pH 6.0, for 15 min at 100 °C. Slides were then transferred to cold PBS with 0.025% (v/v) Tween-20 for 15 min, rinsed in water, followed by quenching of endogenous peroxidases using 3% H2O2 for 10 min. After washing (PBS, 0.025% (v/v) Tween-20, and 1% (w/v) BSA) and blocking, slides were incubated with primary antibodies (anti-OTULIN, Abcam or anti-Ki67, clone SP6, LabVision; see list of antibodies below for further information) at 4 °C overnight. Slides were then washed and incubated with secondary biotinylated antibodies for 30 min at room temperature. Secondary antibodies were labelled using the VECTASTAIN ABC HRP Kit (cat# PK-4001, RRID: AB_2336810, Vector Laboratories) and detected using the DAB (3,3’-diaminobenzidine) Peroxidase (HRP) Substrate Kit (cat# SK-4100, RRID: AB_2336382, Vector Laboratories) according to the manufacturer’s protocols. Slides were counterstained with Mayer’s Haematoxylin, dehydrated, cleared, and mounted with DPX mounting medium. TUNEL (terminal deoxynucleotidyl transferase dUTP nick end labelling) assays were performed on deparaffinised and rehydrated FFPE sections using the ApopTag Peroxidase In Situ Apoptosis Detection kit (cat# S7100, RRID: AB_2661855, Merck Millipore) according to the manufacturer’s instructions and counterstained with Mayer’s Haematoxylin. Slides were dehydrated, cleared, and mounted with DPX mounting medium. All stainings were performed by the Cambridge Stem Cell Institute Histology Core Facility, University of Cambridge, Cambridge, UK.

#### Micrographs and image analysis

Micrographs for analysis of histology, IHC, and TUNEL assays were taken on an Axioplan microscope (Carl Zeiss) mounted with a Leica DFC310 FX camera using the Leica LAS software (version 4.6.0). Contrast, brightness, and colour balance were adjusted using Adobe Photoshop CS6. Counting of stained cells, nuclear diameter measurements, and analysis of fibrotic area were performed in the ImageJ or Fiji software (Schindelin et al., 2012) using the Cell Counter plug-in or Point Tool, the ROI manager, and the Color Threshold function (hue, saturation, and brightness (HSB) colour space), respectively. Scale bars represent 200 μm unless otherwise indicated.

#### Serum analyses

Whole blood from terminal bleeds were collected in 1.5 mL Eppendorf microcentrifuge tubes and left to clot for 1-2 h at room temperature. Serum was collected after centrifugation of the clotted blood for 30 min at 15,000 x*g* and stored at −80 °C until analysis. Serum alpha-Fetoprotein (AFP) concentrations were measured by ELISA using the Mouse alpha-Fetoprotein/AFP Quantikine ELISA Kit (cat# MAFP00; R&D Systems) according to the manufacturer’s instructions. Serum insulin was analysed using the Mouse/Rat Insulin Kit (cat# K152BZC-3; MesoScale Discovery, Rockville, MD) according to the manufacturer’s instructions. Serum levels of albumin, bilirubin, glucose, triglycerides, cholesterol, alanine aminotransferase (ALT), and aspartate aminotransferase (AST) were measured on a fully automated Dimension EXL Analyser (Siemens Healthcare) using the DF13, DF167, DF30, DF69A, DF27, DF143, and DF41A cartridges (Siemens Healthcare), respectively. All analyses were performed at the Core Biochemical Assay Laboratory, Cambridge University Hospitals, Cambridge, UK.

#### Flow cytometry

For analysis of chimerism and the proportion of parental cells remaining in the Control^Chim^ and *Otulin*-KO^Chim^ mice, spleens were dissected and placed in ice-cold PBS containing 3% (v/v) FCS. Single cell suspensions were obtained by gentle mechanical disruption of the tissues using a syringe plunger and a 70 μm cell strainer (Corning). Single cells in a solution of 2% (v/v) FBS in PBS were kept on ice and blocked with anti-CD16/CD32 (Fc*γ*RIIIb/Fc*γ*RIIa) antibodies for 20 min, washed, and then incubated with BrilliantViolet-510-coupled anti-CD45.1 (clone A20, BioLegend) and AlexaFluor-700-coupled anti-CD45.2 (clone 104, eBioscience) antibodies for 30 min before analysis. Fixable Viability Dye eFluor780 (eBioscience) was included to exclude dead cells. Cells were analyzed on a BD Fortessa (BD Bioscience) according to the manufacturer’s standard operating procedures. Data were analysed with FlowJo 10.0.7 (Tree Star). The percentage of parental cells was determined by dividing the number of cells in the CD45.2 (C57BL/6) gate with the sum of cells in the CD45.1 (B6.SJL) and CD45.2 (C57BL/6) gates.

#### Purification of endogenous polyUb conjugates

The GST-tagged TUBE and M1-SUB were expressed and purified from E. coli as previously described (Damgaard et al., 2019). Endogenous polyUb conjugates were purified from mouse livers using the TUBE or M1-SUB affinity reagents as described previously (Damgaard et al., 2019; Fiil et al., 2013; Hrdinka et al., 2016). Briefly, liver tissue was lysed using stainless steel beads (Ø = 5 mm; QIAGEN) for 7 min at 20 Hz on a TissueLyser II (QIAGEN) in buffer containing 20 mM Na2HPO4, 20 mM NaH2PO4, 1% (v/v) NP-40, and 2 mM EDTA, and supplemented with 5 mM N-Ethylmaleimide (NEM) and cOmplete protease inhibitor cocktail (Roche). GST-tagged TUBE (50 μg/ml) or M1-SUB (100 μg/mL) was added directly to the lysis buffer immediately before lysis. Lysates were cleared by centrifugation (16,000 x*g*) at 4 °C for 30 minutes and Glutathione Sepharose 4B resin (GE Healthcare) was added and pulldown performed for 16-20 h at 4 °C on rotation. The resin was washed four times in 1 mL of ice-cold PBS + 0.1% (v/v) Tween-20 (PBS-T) and bound material was eluted by mixing the resin with 1x sample buffer (50 mM Tris pH 6.8, 10% (v/v) glycerol, 100 mM DTT, 2% (w/v) SDS, and 0.01% (w/v) bromophenol blue) and heating to 50 °C for 5 min. Samples were subjected to analysis by immunoblotting.

#### Immunoblotting

Mouse livers were lysed in RIPA buffer (50 mM Tris pH 7.4, 1% NP-40 (v/v), 0.5% deoxycholate (w/v), 0.1% SDS (w/v), 150 mM NaCl, 2 mM EDTA, 5 mM MgCl2) supplemented with cOmplete protease inhibitor cocktail (Roche) and PhosSTOP phosphatase inhibitor (Roche) using stainless steel beads (Ø = 5 mm; QIAGEN) for 7 min at 20 Hz on a TisssueLyser II (QIAGEN). Samples were treated with Benzonase (Novagen, Madison, WI) for 30 min at 4 °C on rotation before lysates were cleared by centrifugation (16,000 x*g*) at 4 °C for 30 minutes. Samples were supplemented with 4x sample buffer to reach a final 1x concentration of 50 mM Tris pH 6.8, 10% (v/v) glycerol, 100 mM DTT, 2% (w/v) SDS, and 0.01% (w/v) bromophenol blue and heated to 95 °C for 5 min. Samples were resolved on 4-12% Bis-Tris NuPAGE or Novex WedgeWell 4-20% Tris-Glycine gels (Life Technologies) and transferred to Trans-Blot Turbo nitrocellulose or polyvinylidene fluoride (PVDF) membranes (Bio-Rad) using the Trans-Blot Turbo transfer system (Bio-Rad). Membranes were blocked in 5% (w/v) skimmed milk powder dissolved in TBS + 0.1% (v/v) Tween-20 (TBS-T) for 30 min and incubated with primary antibodies in TBS-T + 3% (w/v) BSA (Sigma) at 4 °C overnight. Afterwards, immunoblots were washed in TBS-T and incubated at room temperature for 1 h with HRP-coupled secondary antibodies and visualised using Clarity Western or Clarity Max ECL Substrate (Bio-Rad) on a ChemiDoc MP imager (Bio-Rad). Digital images were processed using the Image Lab software (Bio-Rad). Please see below for details on antibodies used in this study. See Supplemental Resources Table for antibodies.

#### Quantitative Real-Time PCR

Total RNA was extracted from mouse liver using the RNeasy Mini Kit (QIAGEN). Liver tissue was lysed in buffer RLT using stainless steel beads (Ø = 5 mm; QIAGEN) for 7 min at 20 Hz on a TissueLyser II (QIAGEN) according to the manufacturer’s instructions. DNase digestion was performed on-column with the RNase-Free DNase Set (QIAGEN) according to the manufacturer’s protocol. Total RNA was reverse transcribed using Quantitect Reverse Transcription Kit (QIAGEN). To ensure complete elimination of genomic DNA, gDNA Wipeout was performed before reverse transcription according to the manufacturer’s instructions. RT-PCR was performed using QuantiFast SYBR Green RT-PCR Kit (QIAGEN) on a Viia7 Real-Time PCR Instrument (Applied Biosystems) with the primers indicated below. Results were normalised to those of 18S rRNA as internal housekeeping control using the 2^(-^*^Δ^*^Ct)^-method. See Table S1 for primer sequences.

#### Nuclei isolation and DNA content analysis

Livers from 8-week old *Otulin*^Δhep^ and control mice, dissected and placed in ice-cold PBS containing 3% (v/v) FCS and stored at −80 °C until their use, were passed through a Falcon 70 μm nylon cell strainer (Corning). Cells were washed in ice-cold Buffer LA (250 mM sucrose, 5 mM MgCl2 and 10 mM Tris-HCl pH 7.4) and pelleted by centrifugation at 600 x*g* in a cold (4 °C) centrifuge twice, discarding the supernatant after each wash. After the second wash, the cell pellets were each resuspended in 1 mL of ice-cold Buffer LB (2 M sucrose, 1 mM MgCl2 and 10 mM Tris-HCl pH 7.4) and centrifuged at 16,000 x*g* for 30 minutes in a cold (4 °C) centrifuge. The supernatants were discarded and the nuclei-containing pellets were kept. This step was repeated three times to obtain clean, white pellets of nuclei, which were resuspended in Buffer LA and stored on ice. For DNA content (ploidy) analysis, nuclei were fixed by drop-wise addition of 5 mL −20 °C 96% ethanol while vortexing and left to fix overnight at −20 °C. Nuclei were pelleted at 300 x*g* in a cold (4 °C) centrifuge and re-suspended in 1 mL of ice-cold PBS containing propidium iodide (40 μg/mL; Sigma) and RNase A (100 μg/mL; Sigma). The samples were incubated on ice for 30 min in the dark and then analysed on an LSRII flow cytometer (BD Bioscience) according to the manufacturer’s standard operating procedures. Data were analysed with FlowJo 10.0.7 (Tree Star).

### QUANTIFICATION AND STATISTICAL ANALYSIS

#### Statistics

Data are presented as individual data points or as means ±SD or SEM as indicated in figure legends. Red bars represent means. Sample number (n) represents the number of independent biological samples in each experiment. Sample numbers and experimental repeats are indicated in figures and figure legends. Data were analysed using the unpaired, two-sided Student’s *t* test of the null hypothesis as indicated. Samples sizes were estimated based on pilot experiments. Differences in means were considered statistically significant at P<0.05. Significance levels are: * P<0.05; ** P<0.01; *** P<0.001; **** P<0.0001; n.s., non-significant. Analyses were performed using the GraphPad Prism software version 7.0b.

### DATA AND CODE AVAILABILITY

This study did not generate any new original code or datasets for analysis.

