## Supplemental Figures and Information for "OTULIN protects the liver against cell death, inflammation, fibrosis, and cancer"

### SUPPLEMENTAL INFORMATION

**Supplemental Figures S1-7**
**Supplemental Table S1**
**Supplemental Recourses Table**
**Supplemental Legends**

### Supplementary Figure S1

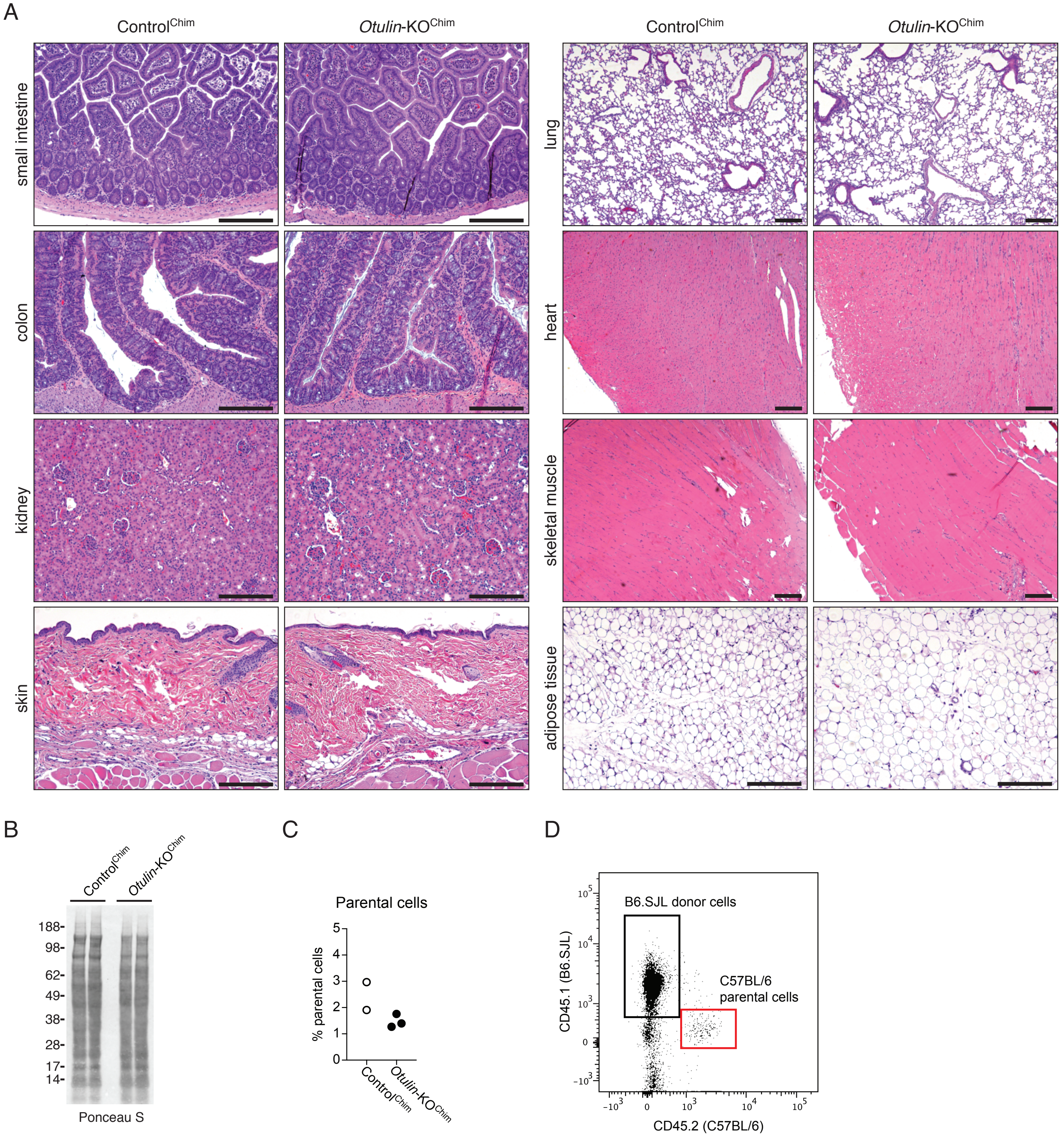

### Supplementary Figure S2

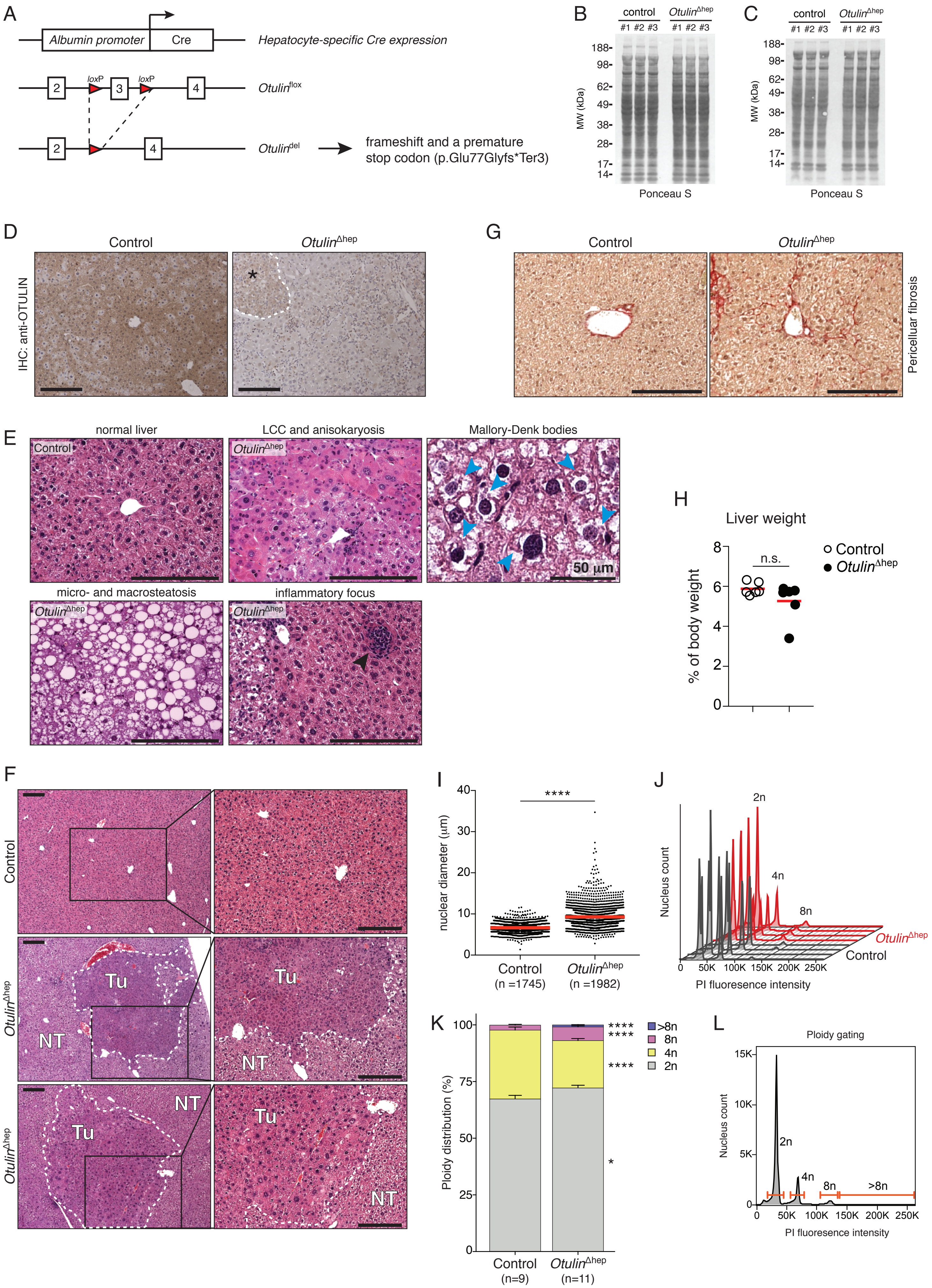

### Supplementary Figure S3

A

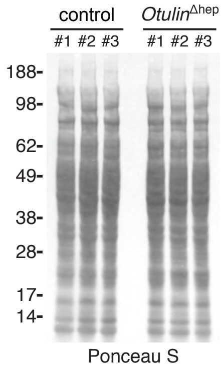


B

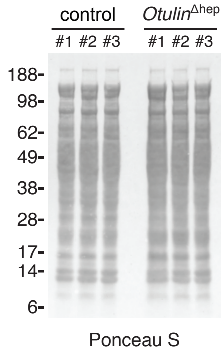

### Supplementary Figure S4

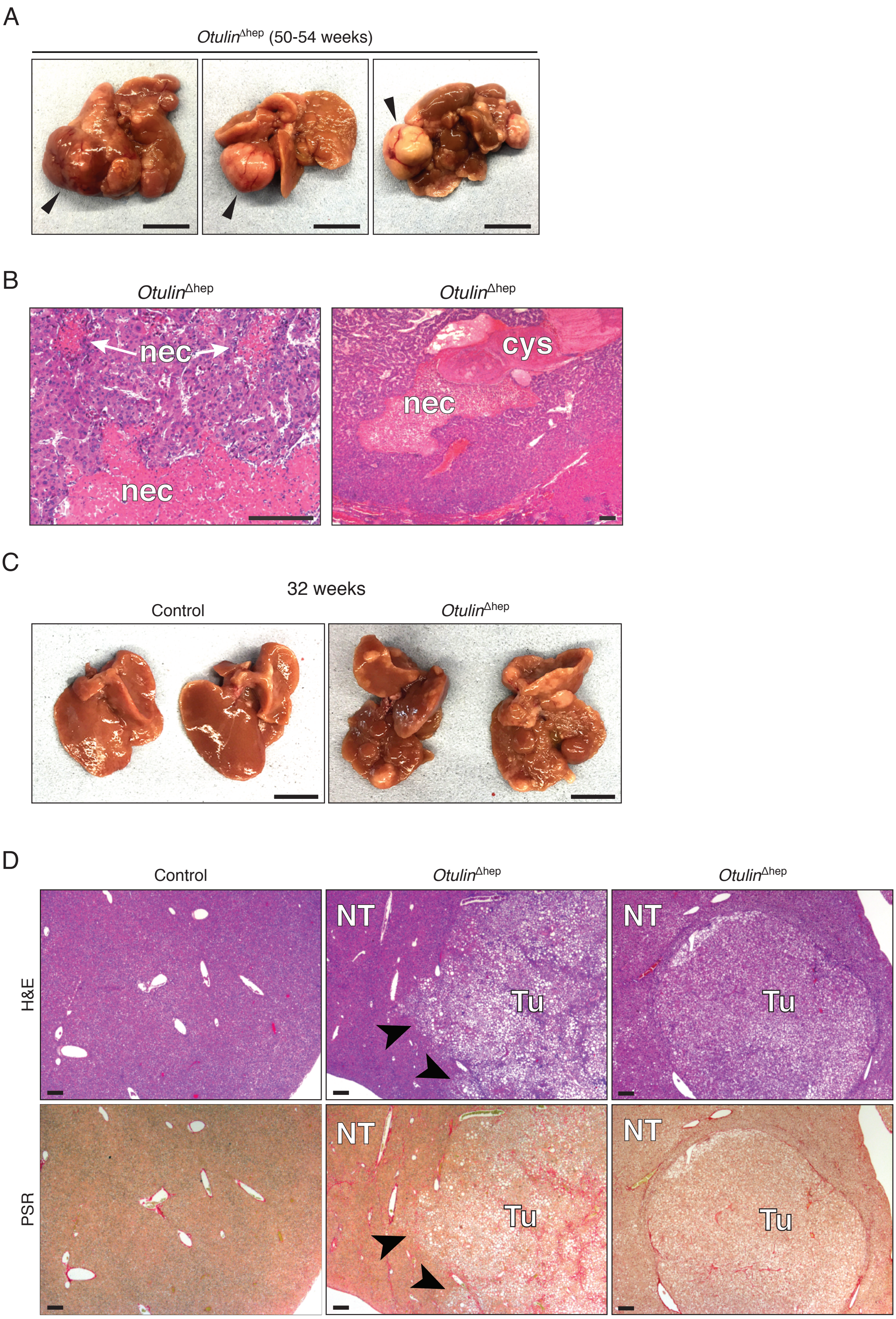

### Supplementary Figure S5

A

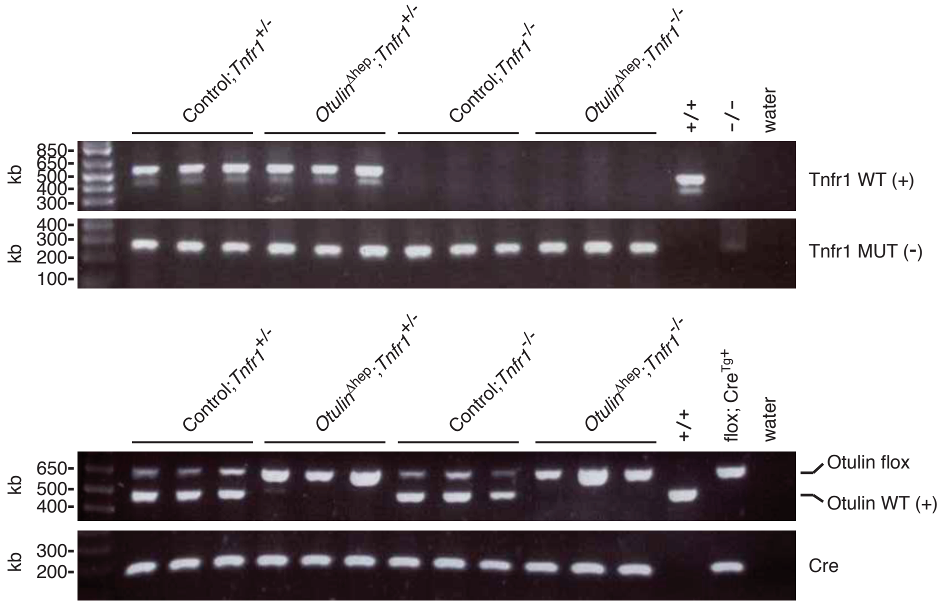


B

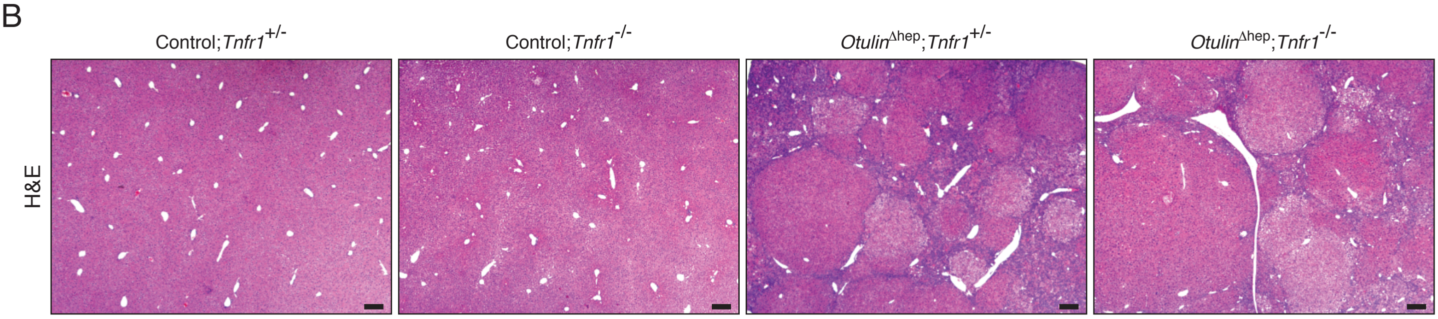

**Supplementary Figure S6**

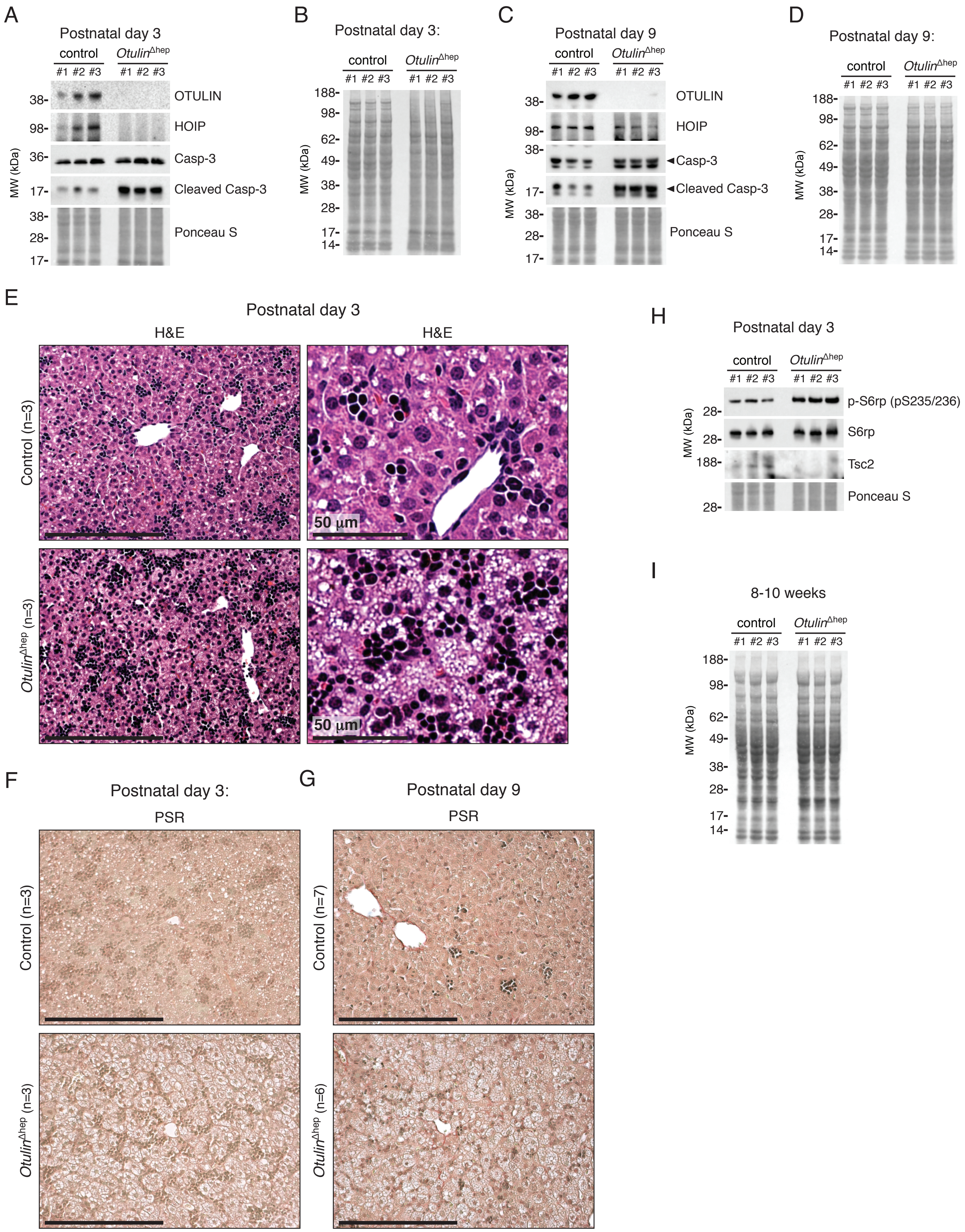

### Supplementary Figure S7

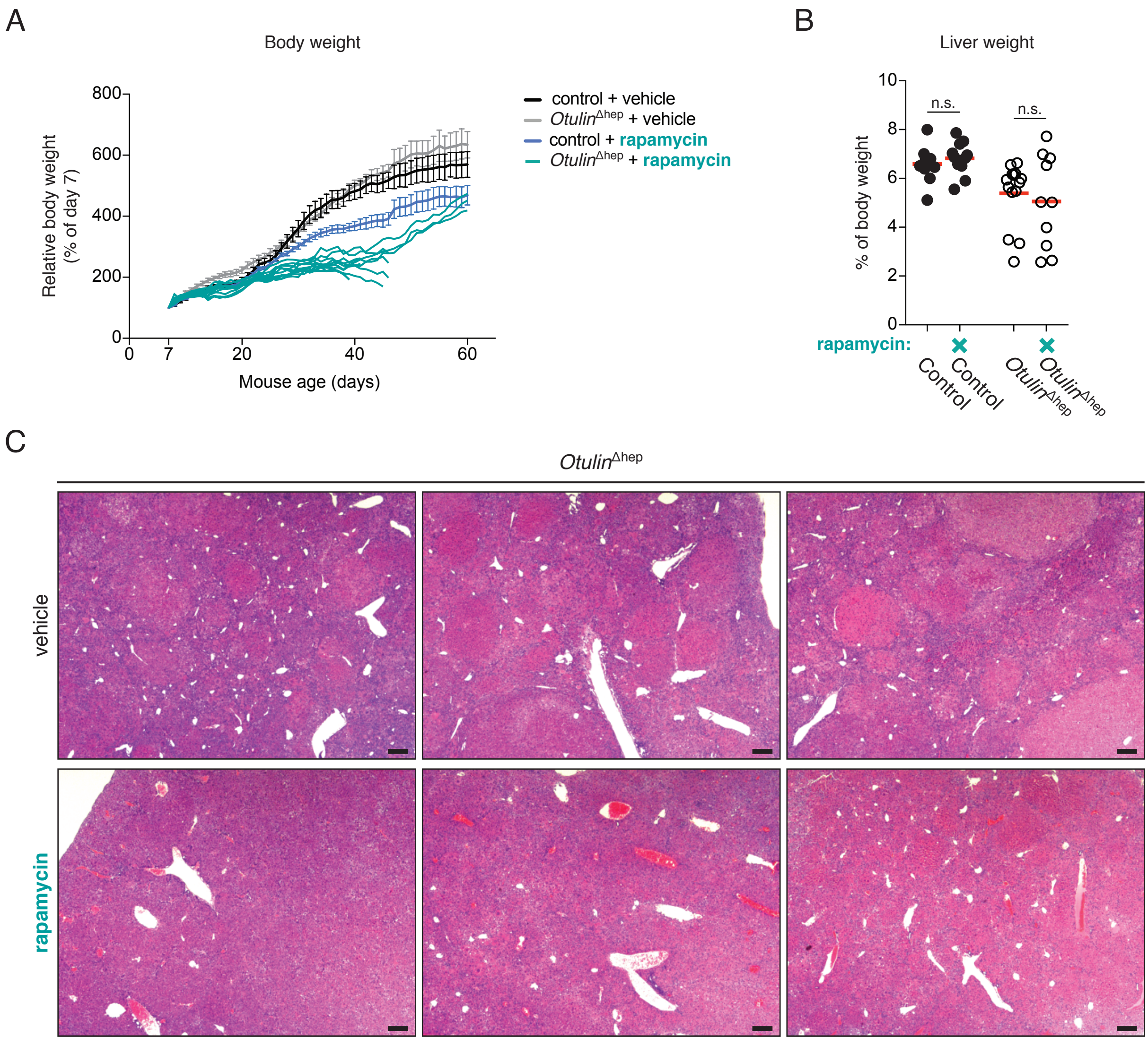

### Table S1: primers for RT-PCR

| Target | Forward primer | Reverse primer |
|---|---|---|
| *18S rRNA* | 5′-GTAACCCGTTGAACCCCATT-3′ | 5′-CCATCCAATCGGTAGTAGCG-3′ |
| *Tnf* | 5′-CCACCACGCTCTTCTGTCTAC-3′ | 5′-AGGGTCTGGGCCATAGAACT-3′ |
| *Il6* | 5′-TAGTCCTTCCTACCCCAATTTCC-3′ | 5′-TTGGTCCTTAGCCACTCCTTC-3′ |
| *Il1b* | 5′-CAATGGACAGAATATCAAC-3′ | 5′-ACAGGACAGGTATAGATT-3′ |
| *Tnfaip3 (A20)* | 5′-TTCCTCAGGACCAGGTCAGT-3′ | 5′-AAGCTCGTGGCTCTGAAAAC-3′ |
| *Cd68* | 5′-TGTCTGATCTTGCTAGGACCG-3′ | 5′-GAGAGTAACGGCCTTTTTGTGA-3′ |
| *Acta2 (Smooth muscle actin)* | 5′-CCCCTGAAGAGCATCGGACA-3′ | 5′-TGGCGGGGACATTGAAGGT-3′ |
| *Ccnd1 (Cyclin D1)* | 5′-GCCGAGAAGTTGTGCATCTAC-3′ | 5′-GGAGAGGAAGTGTTCGATGAA-3′ |
| *Ctgf* | 5′-GCCCTAGCTGCCTACCGACT-3′ | 5′-GCCCATCCCACAGGTCTTAGA-3′ |
| *Gpc3* | 5′-CTGAGCCGGTGGTTAGCC-3′ | 5′-TCACTTTCACCATCCCGTCA-3′ |
| *Igf2* | 5′-ACATGCTGCCCAAGTAACC-3′ | 5′-CTGACAAAGATGGCCCATAG-3′ |
| *Afp* | 5′-CTCAGCGAGGAGAAATGGTC-3′ | 5′-GAGTTCACAGGGCTTGCTTC-3′ |
| *H19* | 5′-CAGGGCTAGTCCGCTCAA-3′ | 5′-AACAGACGGCTTCTACGACAA-3′ |
| *Klf4* | 5′-CGGACCACCTTGCCTTACACA-3′ | 5′-TGACTTGCTGGGAACTTGACC-3′ |
| *Aldh1 (Aldh17a)* | 5′-GGTGAACATTGTCCCTGGTTAT-3′ | 5′-GACACTTTGTCGATGTCCATGT-3′ |
| *Cd133 (Prom1)* | 5′-TGGAGCTACCTGCGGTTTAGA-3′ | 5′-GGACCTGTGATTGCGATAATGA-3′ |

**SUPPLEMENTAL RESOURCES TABLE**

| REAGENT or RESOURCE | SOURCE | IDENTIFIER |
|---|---|---|
| **Antibodies** | | |
| Rabbit polyclonal anti-OTULIN | Abcam | Cat#ab151117; RRID: AB_2728115 |
| Rabbit polyclonal anti-mouse HOIP | Tokunaga et al., 2011 | N/A |
| Mouse monoclonal anti-HOIL-1/RBCK1 (clone 2E2) | Merck Millipore | Cat#MABC576; RRID: AB_2737058 |
| Rabbit polyclonal anti-SHARPIN | ProteinTech | Cat#14626-1-AP; RRID: AB_2187734 |
| Rabbit monoclonal anti-CYLD (clone D1A10) | Cell Signaling Technology | Cat#8462; RRID: AB_10949157 |
| Rabbit polyclonal anti-IκBα | Cell Signaling Technology | Cat#9242; RRID: AB_10694550 |
| Rabbit monoclonal anti-p65/RelA (clone D14E12) | Cell Signaling Technology | Cat#8242; RRID: AB_10859369 |
| Rabbit monoclonal phospho-p65/RelA (S563) (clone 93H1) | Cell Signaling Technology | Cat#3033; RRID: AB_331285 |
| Rabbit monoclonal anti-ERK1/2 (clone 137F5) | Cell Signaling Technology | Cat#4695; RRID: AB_390779 |
| Rabbit monoclonal anti-phospho-ERK1/2 (T202/Y204) (clone D13.14.E4) | Cell Signaling Technology | Cat#4370; RRID: AB_2315112 |
| Mouse monoclonal anti-p38 (clone M138) | Abcam | Cat#ab31828; RRID: AB_881839 |
| Rabbit monoclonal anti-phospho-p38 (T180/Y182) (clone ERP18120) | Abcam | Cat#ab195049; RRID: AB_2576214 |
| Rabbit monoclonal anti-Caspase-3 (clone D3R6Y) | Cell Signaling Technology | Cat#14220; RRID: AB_2798429 |
| Rabbit monoclonal anti-cleaved Caspase-3 (D175) (clone 5A1E) | Cell Signaling Technology | Cat#9664; RRID: AB_2070042 |
| Rabbit monoclonal anti-S6rp (clone 5G10) | Cell Signaling Technology | Cat#2217; RRID: AB_331355 |
| Rabbit monoclonal anti-phospho-S6rp (S235/S236) (clone D57.2.2E) | Cell Signaling Technology | Cat#4858; RRID: AB_916156 |
| Rabbit monoclonal anti-TSC1/Hamartin (clone D43E2) | Cell Signaling Technology | Cat#6935; RRID: AB_10860420 |
| Rabbit monoclonal anti-TSC2/Tuberin (clone D57A9) | Cell Signaling Technology | Cat#3990; RRID: AB_2209986 |
| Rabbit monoclonal anti-Rheb (clone E1G1R) | Cell Signaling Technology | Cat#13879; RRID: AB_2721022 |
| Rabbit polyclonal anti-mTOR | Cell Signaling Technology | Cat#2972; RRID: AB_330978 |
| Rabbit polyclonal anti-phospho-mTOR (S2448) | Cell Signaling Technology | Cat#2971; RRID: AB_330970 |
| Mouse monoclonal anti-Ubiquitin (clone Ubi-1) | Novus Biologicals | Cat#NB300-130; RRID: AB_2238516 |
| Rabbit monoclonal anti-linear Ubiquitin (M1-polyUb) (clone 1E3) | Merck Millipore | Cat# MABS199; RRID: AB_2576212 |
| Rabbit monoclonal anti-Ki67 (clone SP6) | LabVision | Cat#RM-9106-R7; RRID: AB_149920 |
| Rat monoclonal anti-mouse CD16/CD32 (clone 2.4G2) | BioXCell | Cat#BE0307; RRID: AB_2736987 |
| Mouse monoclonal anti-mouse CD45.1 BrilliantViolet-510-coupled (clone A20) | BioLegend | Cat#110741; RRID: AB_2563378 |
| Mouse monoclonal anti-mouse CD45.2 AlexaFluor-700-coupled (clone 104) | eBioscience | Cat#56-0454-82; RRID: AB_657752 |
| Donkey polyclonal anti-rabbit IgG HRP-coupled | GE Healthcare | Cat#NA934; RRID: AB_772206 |

| | | |
|---|---|---|
| Sheep polyclonal anti-mouse IgG HRP-coupled | GE Healthcare | Cat#NXA931; RRID: AB_772209 |
| **Chemicals, Peptides, and Recombinant Proteins** | | |
| Rapamycin | LC Laboratories | Cat#R-5000; CAS: 53123-88-9 |
| GST-tagged TUBE | Laboratory stock | Hrdinka et al., 2016 |
| GST-tagged M1-SUB | Laboratory stock | Fiil et al., 2013 |
| **Critical Commercial Assays** | | |
| ApopTag Peroxidase In Situ Apoptosis Detection kit (TUNEL assay) | Merck Millipore | Cat#S7100; RRID: AB_2661855 |
| Mouse alpha-Fetoprotein/AFP Quantikine ELISA Kit | R&D Systems | Cat#MAFP00 |
| Mouse/Rat Insulin Kit | MesoScale Discovery | Cat#K152BZC-3 |
| RNeasy Mini Kit | QIAGEN | Cat#74104 |
| RNase-Free DNase Set | QIAGEN | Cat#79254 |
| Quantitect Reverse Transcription Kit | QIAGEN | Cat#205313 |
| QuantiFast SYBR Green RT-PCR Kit | QIAGEN | Cat#205314 |
| **Experimental Models: Organisms/Strains** | | |
| Mouse: C57BL/6-*Otulin*$^{tm1Anjm}$ | Laboratory of ANJ McKenzie | Damgaard et al, 2016 |
| Mouse: C57BL/6-*Otulin*$^{tm1Anjm}$-Gt(ROSA)26SorCreERT2 | Laboratory of ANJ McKenzie | Damgaard et al, 2016 |
| Mouse: C57BL/6-*Speer6-ps1*$^{Tg(Alb-cre)21Mgn}$ (Alb-Cre) | JAX | Cat#003574; RRID: IMSR_JAX:003574 |
| Mouse: C57BL/6-*Tnfrsf1a*$^{tm1Imx}$ (TNFR1-/-) | JAX | Cat#003242; RRID: IMSR_JAX:003242 |
| **Oligonucleotides** | | |
| Primers for RT PCR analyses, see Table S1 | This paper | N/A |
| **Software and Algorithms** | | |
| Fiji / ImageJ | https://fiji.sc/ | Schindelin et al., 2012 |
| **Other** | | |
| Glutathione Sepharose 4B resin | GE Healthcare | Cat#17-0756-01 |
| AML Laboratories (histology services) | http://www.amllabs.com/ | N/A |
| Cambridge University Hospitals' Core Biochemical Assay Laboratory (clinical biochemical analyses) | https://www.cuh.nhs.uk/core-biochemical-assay-laboratory | N/A |

### Supplemental Legends

**Figure S1. Analysis of Control$^{Chim}$ and *Otulin*-KO$^{Chim}$ mice. Related to Figure 1.**

**(A)** Micrographs of H&E stained tissue sections from Control$^{Chim}$ and *Otulin*-KO$^{Chim}$ mice at the end of the experiment shown in Figure 1B. Micrographs are representative of two mice in each group.

**(B)** Uncropped Ponceau S stained membrane from Figure 1G.

**(C)** Percentage of parental (CD45.2+) cells in the spleens of Control$^{Chim}$ and *Otulin*-KO$^{Chim}$ at the end of the experiment shown in Figure 1B.

**(D)** Representative dot plot from flow cytometric analysis of parental (CD45.2+) and B6.SLJ (CD45.1+) splenocytes used to generate the plot in (C).

**Figure S2. Generation and analysis of mice with hepatocyte-specific deletion of *Otulin* (*Otulin*$^{\Delta hep}$ mice). Related to Figure 2.**

**(A)** Schematic showing the strategy used to generate mice with hepatocyte-specific deletion of *Otulin*. Numbers denote *Otulin* exons.

**(B)** Uncropped Ponceau S stained membrane from Figure 2B.

**(C)** Uncropped Ponceau S stained membrane from Figure 2C.

**(D)** Immunohistochemical analysis of OTULIN expression in *Otulin*$^{\Delta hep}$ and control mice at the age of 8-10 weeks shows clonal areas of hepatocytes that retain OTULIN protein expression (asterisk), likely due to incomplete penetrance and recombination efficiency of the *Alb*-Cre transgene. Micrographs are representative of two mice of each genotype.

**(E)** Micrographs of H&E stained liver sections from *Otulin*$^{\Delta hep}$ and control mice at the age of 8-10 weeks show histological features present in the diseased livers of *Otulin*$^{\Delta hep}$ mice. Black arrowhead indicates inflammatory focus. Blue arrowheads indicate Mallory-Denk bodies. LCC, large cell change. Micrographs are representative of six mice of each genotype.

**(F)** Micrographs of H&E stained liver sections from *Otulin*$^{\Delta hep}$ and control mice at the age of 8-10 weeks show formation of pre-malignant tumours (dysplastic nodules) in *Otulin*$^{\Delta hep}$ livers. Micrographs are representative of six mice of each genotype. Tu, tumour. NT, non-tumour.

**(G)** High magnification micrographs of PSR stained liver sections (shown in Figure 2E) from *Otulin*$^{\Delta hep}$ and control mice aged 8-10 weeks showing pericellular collagen deposition.

**(H)** Relative liver weights from *Otulin*$^{\Delta hep}$ (n=6) and control mice (n=6) at the age of 8-10 weeks. Each data point represents one mouse. Red bars indicate means. Data were analysed using the unpaired, two-sided Student's *t* test. n.s., non-significant.

**(I)** Quantification of nuclear diameter of hepatocytes from *Otulin*$^{\Delta hep}$ and control mice at the age of 8-10 weeks. Two fields of view were quantified from each of six *Otulin*$^{\Delta hep}$ mice and six control mice. Each data point represents one mouse. Red bars indicate means. Data were analysed using an unpaired, two-sided Student's *t* test. n.s., non-significant.

**(J)** Representative staggered histograms from flow cytometric analysis of DNA content in nuclei isolated from *Otulin*$^{\Delta hep}$ and control mice at the age of 8-10 weeks showing increased proportions of nuclei with DNA content ≥8n.

**(K)** Quantification of flow cytometric analysis as shown in (I). Data represent mean +SEM. Data were analysed using the unpaired, two-sided Student's *t* test.

**(L)** Gating strategy on a representative liver sample for flow cytometric analysis as shown in (I-J).

**Figure S3. Analysis of liver disease in *Otulin*$^{\Delta hep}$ mice. Related to Figure 3.**

**(A)** Uncropped Ponceau S stained membrane from Figure 3F.

**(B)** Uncropped Ponceau S stained membrane from Figure 3H.

**Figure S4. Analysis of hepatocellular carcinoma in *Otulin*$^{\Delta hep}$ mice. Related to Figure 4.**

**(A)** Representative macroscopic appearance of *Otulin*$^{\Delta hep}$ livers at the age of 50-54 weeks. Arrowheads indicate highly vascularised tumours. Scale bars indicate 1 cm.

**(B)** Micrographs of H&E stained tumours from *Otulin*$^{\Delta hep}$ mice aged 50-54 weeks. nec, necrotic area. cys, cystic lesion.

**(C)** Representative macroscopic appearance of *Otulin*$^{\Delta hep}$ and control livers at the age of 32 weeks. Scale bars indicate 1 cm.

**(D)** Micrographs of H&E (top panels) and PSR (bottom panels) stained liver sections from *Otulin*$^{\Delta hep}$ (n=5) and control mice (n=8) at the age of 32 weeks. Arrowheads indicate areas of poor tumour demarcation. Tu, tumour. NT, non-tumour.

**Figure S5. Analysis of OTULIN and TNFR1 double-deficient livers. Related to Figure 5.**

**(A)** PCR genotyping of *Otulin*$^{\Delta hep}$ mice, *Otulin*$^{\Delta hep}$;*Tnfr1*$^{-/-}$, and their respective controls show co-deletion of *Otulin* and *Tnfr1* as expected. kb, kilobases.

**(B)** Micrographs of H&E stained liver sections from *Otulin*$^{\Delta hep}$ mice, *Otulin*$^{\Delta hep}$;*Tnfr1*$^{-/-}$ mice, and their respective controls at the age of 8-12 weeks showing no difference in histopathological changes between *Otulin*$^{\Delta hep}$ mice and *Otulin*$^{\Delta hep}$;*Tnfr1*$^{-/-}$ mice.

**Figure S6. Analysis of neonatal *Otulin*$^{\Delta hep}$ and control mice. Related to Figure 6.**

**(A)** Immunoblot analysis of OTULIN, HOIP, and caspase-3 in whole-liver lysates from three *Otulin*$^{\Delta hep}$ and three control mice aged 3 days.

**(B)** Uncropped Ponceau S stained membrane from Figures S6A and S6H.

**(C)** Immunoblot analysis of OTULIN, HOIP, and caspase-3 in whole-liver lysates from three *Otulin*$^{\Delta hep}$ and three control mice aged 9 days.

**(D)** Uncropped Ponceau S stained membrane from Figure S6C.

**(E)** Micrographs of H&E stained liver sections from *Otulin*$^{\Delta hep}$ and control mice at the age of 3 days (P3).

**(F-G)** Micrographs of PSR stained liver sections from *Otulin*$^{\Delta hep}$ and control mice at the age of 3 days (P3) (F) and 9 days (P9) (G).

**(H)** Immunoblot analysis of mTOR pathway components and activation in whole-liver lysate from three *Otulin*$^{\Delta hep}$ mice and three controls aged 3 days.

**(I)** Uncropped Ponceau S stained membrane from Figure 6I.

**Figure S7. Analysis of rapamycin-treated *Otulin*$^{\Delta hep}$ and control mice. Related to Figure 7.**

**(A)** Relative body weight for *Otulin*$^{\Delta hep}$ and control mice treated with rapamycin or vehicle as indicated. Each rapamycin-treated *Otulin*$^{\Delta hep}$ mouse is represented by an individual (cyan) line. The mean weights (±SEM) are shown for the other experimental groups. Data were pooled from two independent experiments.

**(B)** Relative liver weights from *Otulin*$^{\Delta hep}$ and control mice at the age of 6 weeks treated with rapamycin or vehicle as indicated. Each data point represents one mouse. Red bars indicate means. Data were analysed using the unpaired, two-sided Student's *t* test. n.s., non-significant.

**(C)** Representative micrographs of H&E stained liver sections from three vehicle-treated and three rapamycin-treated *Otulin*$^{\Delta hep}$ mice at the age of 6 weeks.

**Table S1. RT-PCR primers. Related to STAR Methods.**

Oligonucleotide sequences of primers used for RT-PCR analyses.
